# Discovery of a carbonic anhydrase-Rubisco complex within the alpha-carboxysome

**DOI:** 10.1101/2021.11.05.467472

**Authors:** Cecilia Blikstad, Eli J. Dugan, Thomas G. Laughlin, Julia B. Turnšek, Mira D. Liu, Sophie R. Shoemaker, Nikoleta Vogiatzi, Jonathan P. Remis, David F. Savage

## Abstract

Carboxysomes are proteinaceous organelles that encapsulate key enzymes of CO_2_ fixation - Rubisco and carbonic anhydrase - and are the centerpiece of the bacterial CO_2_ concentrating mechanism (CCM). In the CCM, actively accumulated cytosolic bicarbonate diffuses into the carboxysome and is converted to CO_2_ by carbonic anhydrase, producing a high CO_2_ concentration near Rubisco and ensuring efficient carboxylation. Self-assembly of the α-carboxysome is orchestrated by the intrinsically disordered scaffolding protein, CsoS2, which interacts with both Rubisco and carboxysomal shell proteins, but it is unknown how the carbonic anhydrase, CsoSCA, is incorporated into the α-carboxysome. Here, we present the structural basis of carbonic anhydrase encapsulation into α-carboxysomes from *Halothiobacillus neapolitanus*. We find that CsoSCA interacts directly with Rubisco via an intrinsically disordered N-terminal domain. A 1.98 Å single-particle cryo-electron microscopy structure of Rubisco in complex with this peptide reveals that CsoSCA binding is predominantly mediated by a network of hydrogen bonds. CsoSCA’s binding site overlaps with that of CsoS2 but the two proteins utilize substantially different motifs and modes of binding, revealing a plasticity of the Rubisco binding site. Our results advance the understanding of carboxysome biogenesis and highlight the importance of Rubisco, not only as an enzyme, but also as a central hub for mediating assembly through protein interactions.

## Introduction

Carbonic anhydrase (CA) catalyzes the rapid interconversion between carbon dioxide (CO_2_) and bicarbonate (HCO_3_^-^). Due to the centrality of this reaction in metabolism, CA is an essential protein in all organisms where it has been tested (1–3). In photosynthesis, CA’s role is often to supply the enzyme Ribulose-1,5-bisphosphate carboxylase-oxygenase (Rubisco) - the carboxylase of the Calvin-Benson-Bassham cycle - with CO_2_ to ensure fast fixation (3). Rubisco has modest turnover numbers and fails to distinguish between CO_2_ and the competing off-target substrate of O_2_ (4–6). To overcome Rubisco’s limitation, plants, algae and some bacteria have evolved different types of CO_2_ Concentrating Mechanisms (CCMs) which concentrate CO_2_ near Rubisco (7). This ensures saturation of Rubisco’s active sites with CO_2_, competitive inhibition of oxygenation, and an increase in overall carbon assimilation rates. Importantly, to understand the role of a CA in a CCM, it is essential to understand enzyme localization and regulation (3).

The bacterial CCM is present in all cyanobacteria and many proteobacteria and consists of two main components: (I) energy-coupled inorganic carbon transporters that actively accumulate bicarbonate in the cytosol and (II) a proteinaceous bacterial organelle called the carboxysome, which co-encapsulates Rubisco and CA within a capsid-like protein shell (8–11). The accumulated HCO_3_^-^ diffuses into the carboxysome where it is rapidly converted to CO_2_ by CA while diffusion out of the structure is likely restricted by the shell (12, 13). This produces a locally high CO_2_ concentration within the carboxysome and enables efficient Rubisco carboxylation (14).

Microbiology and biochemistry shows there are, in fact, two types of carboxysomes which have evolved convergently (8, 15). These are the α-type found in oceanic cyanobacteria and proteobacteria (Fig. 1A) and the β-type found in freshwater cyanobacteria. In both instances, CA localization in the carboxysome is essential for growth in present day atmospheric CO_2_ concentrations (16–18). In contrast, CA activity in the cytosol has been shown to short-circuit the CCM leading to high CO_2_ requiring phenotypes (19). Efficient encapsulation and regulation of CA activity is thus crucial for cell survival. All α-carboxysomes contain a β-CA, CsoSCA (20–23), while β-carboxysomes have either an active γ-CA domain on the scaffolding protein CcmM (24, 25) or a β-CA named CcaA (26). While the mechanism of CA incorporation into the β-carboxysome is understood (27, 28), it is unknown how CsoSCA incorporates into the α-carboxysome.

**Figure 1:**
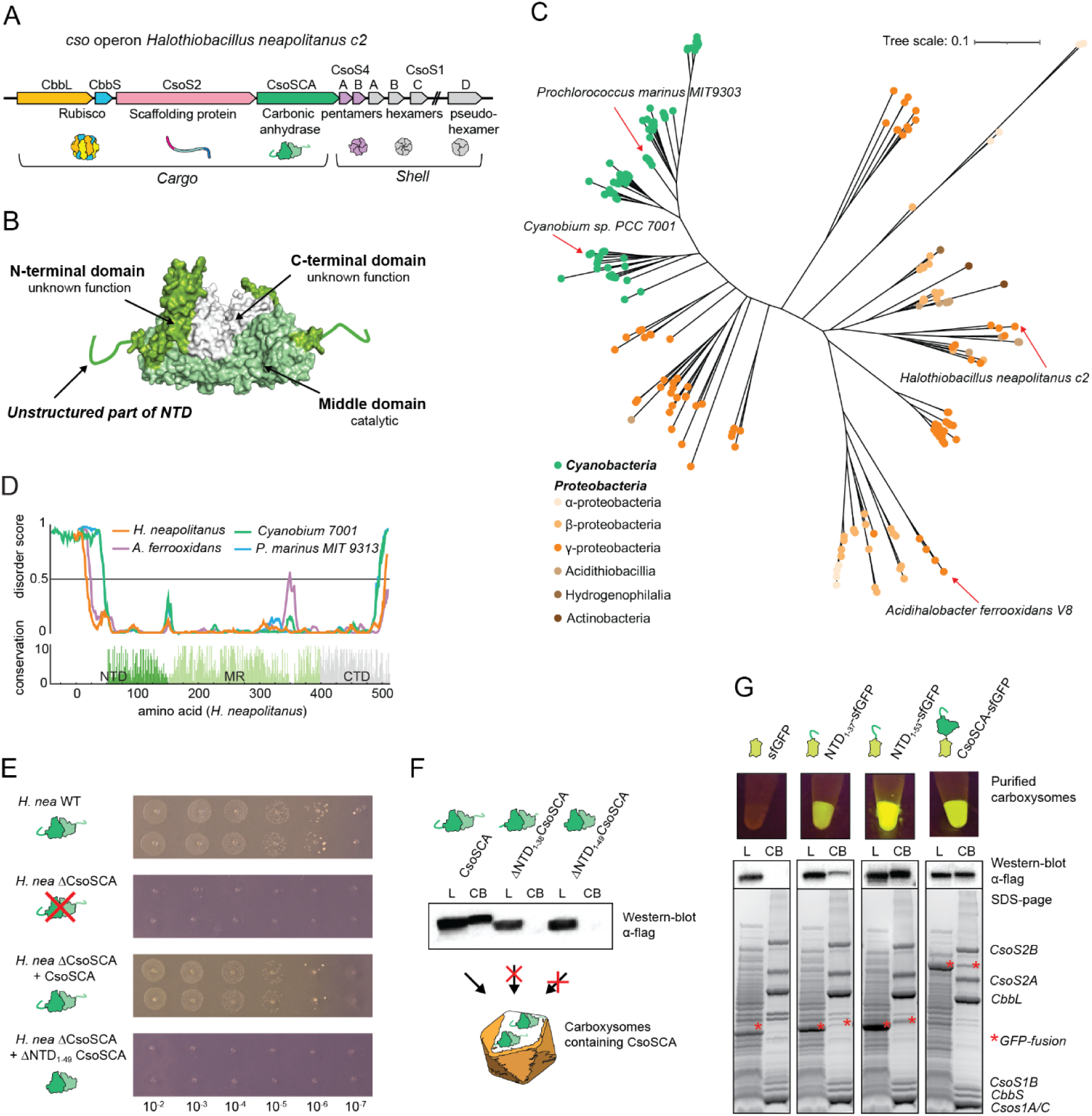
An intrinsically disordered, poorly conserved N-terminal peptide is essential and sufficient for CsoSCA encapsulation. (A) Schematic of the *cso* operon (carboxysome operon) in *H. neapolitanus.* The 10 gene set consists of Rubisco large and small subunit, the scaffolding protein CsoS2, the carbonic anhydrase CsoSCA and six shell proteins (CsoS4A/B, CsoS1A/B/C and CsoSD1). In the native organism, CsoS1D is transcribed from an adjacent locus while in the synthetic pHnCB10 plasmid, all genes are in a single operon. (B) Surface representation structure of CsoSCA dimer from *H. neapolitanus* (pdb: 2FGY). N-terminal domain (dark green) consists of a ∼50 aa long unstructured peptide followed by a folded ɑ-helical domain with unknown function. Middle domain (light green) contains the active site. The C-terminal domain (white) appears to be a gene duplication of the catalytic domain but lacks essential active site residues. (C) Maximum-likelihood phylogenetic tree of CsoSCA. Cyanobacterial homologs are colored in green and proteobacteria homologous in an orange/brown gradient. Scale bar, 0.1 substitutions per site. (D) Disorder score of four representative CsoSCA homologous calculated using Disopred3 and conservation calculated from multiple sequence alignment. (E) Complementation of full-length *csoSCA* rescues growth of a *csoSCA* knock-out in *H. neapolitanus* while complementation with a NTD truncated variant, *ΔNTD_1-49_CsoSCA*, fails to rescue growth. (F) Western-blot analysis detecting C-terminally flag-tagged CsoSCA in lysate (L) and enriched carboxysomes (CB) fractions of carboxysomes produced heterologously in *E. coli*. Synthetic carboxysomes consist of the full cso operon (Fig. 1A), with either wild-type CsoSCA or a N-terminal truncated variant (ΔNTD_1-37_CsoSCA or ΔNTD_1-49_CsoSCA). CsoSCA is not detected in carboxysomes with N-terminal truncated CsoSCA variants. (G) Fusing the unstructured NTD of CsoSCA (37 or 53 residues) to sfGFP targets the fusion protein to synthetic carboxysomes produced heterologously in *E. coli* while untagged sfGFP does not target to carboxysomes. The control with full-length CsoSCA-sfGFP also produces fluorescent carboxysomes. Panel shows fluorescence of purified carboxysomes, western-blot analysis against flag-tagged sfGFP and SDS-PAGE of lysate (L) and purified carboxysomes (CB). In (F) and (G): The L sample contains detergent for lysing the cells (B-PER II) resulting in a small band shift on the SDS-page, explaining the slightly lower CsoSCA and NTD_1-37_/NTD_1-53_ band in L compared to CB.

CsoSCA belongs to its own subclass of β-CAs and uniquely consists of three domains: an N-terminal domain, a middle/catalytic domain and a C-terminal domain (Fig. 1B) (22). X-ray structural analysis has shown that the catalytic domain contains the zinc binding site as well as catalytic residues essential for CA activity. The C-terminal domain appears to be an ancient gene duplication of the catalytic domain but lacks the zinc binding residues. The N-terminal domain (NTD) consists of an unstructured N-terminal peptide followed by a ∼100 residue α-helical domain, which lacks homology to any other known protein. The function of this domain is mysterious and has been speculated to be involved in the encapsulation process (22).

Here, we used biolayer interferometry (BLI) to screen CsoSCA for binding to all α-carboxysome proteins and identified Rubisco as its interaction partner. We show that the Rubisco interaction and encapsulation into carboxysomes is dependent on CsoSCA’s unique intrinsically disordered N-terminal peptide. Using this peptide, we targeted foreign cargo into the carboxysome, demonstrating that this sequence is sufficient for encapsulation. We further determined a 1.98 Å single-particle cryo-electron microscopy (cryo-EM) structure of Rubisco in complex with the NTD peptide. The structure reveals that the peptide interacts with Rubisco at a site overlapping with a recently identified site responsible for targeting Rubisco to the α-carboxysome via interaction with the carboxysomal scaffolding protein CsoS2 (29). Thus, our work identifies a previously unknown supercomplex found inside the α-carboxysome and highlights a surprising flexibility in the scope of protein-protein interactions which lead to α-carboxysome self-assembly.

## Results

### CsoSCA’s N-terminus is necessary and sufficient for encapsulation

In order to identify putative mechanisms for encapsulation of CsoSCA into the carboxysome, we first started with a bioinformatic examination of the CsoSCA protein. Phylogenetic analysis revealed that CsoSCA from cyanobacteria and proteobacteria cluster into two separate subfamilies (Fig. 1C, SI Fig. 1 and SI Table 1). The cyanobacterial subfamily divides into two clusters. The proteobacteria subfamily is more diverse but has three distinct clusters, including a transition cluster more closely related to the cyanobacterial subfamily. Multiple sequence alignment (MSA) and calculated conservation score reveal a poorly conserved N-terminal region of the NTD while the rest of the protein is highly conserved (Fig. 1D). In our model organism, the γ-proteobacterium *H. neapolitanus,* the sequence conservation of CsoSCA starts with residue H51. Using representatives from the different clusters of the phylogenetic tree, it was revealed that this ∼30-130 amino acid long NTD is predicted to be disordered (Fig. 1D). Even though the NTD primary sequence is not conserved, its existence among all species is suggestive of a function.

**Table 1:**
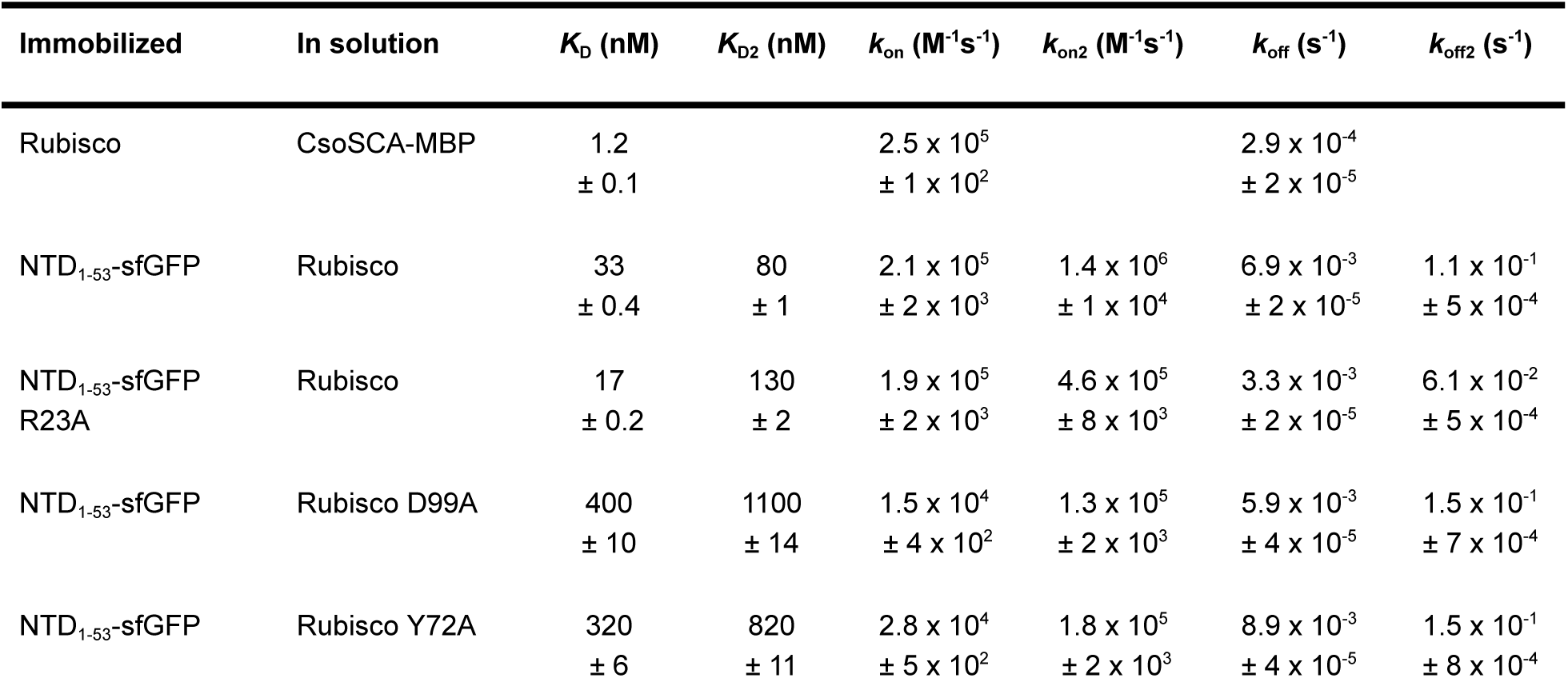
Binding constants and kinetics for CsoSCA and NTD_1-53_-sfGFP binding to Rubisco measured with BLI.

To investigate the role of CsoSCA’s unstructured NTD-peptide, a *csoSCA* deletion strain of *H. neapolitanus* (*ΔCsoSCA*) was complemented with a version of CsoSCA lacking its first 49 residues (*ΔNTD_1-49_CsoSCA)*. The *ΔNTD_1-49_CsoSCA* strain failed to grow in air (Fig. 1E) whereas complementation with full-length CsoSCA rescued growth, indicating an essential role for CsoSCA’s NTD. Synthetic carboxysome operons containing truncated CsoSCAs (*ΔNTD_1-37_CsoSCA* and *ΔNTD_1-49_CsoSCA*) were heterologously expressed in *E. coli*. Western blot analysis of enriched carboxysome (Fig. 1F) fractions showed that neither of these constructs produced carboxysomes containing CsoSCA, suggesting that the growth defect seen in the phenotyping experiment is due to an inability to encapsulate CsoSCA into the carboxysome when the N-terminal peptide (NTD_1-49_-peptide) is removed.

To further test the role of the CsoSCA-NTD in encapsulation, we sought to target foreign cargo into the carboxysome via fusion with peptides derived from the NTD. Monomeric superfolder GFP (sfGFP) fused with either the first 37 (NTD_1-37_-sfGFP) or first 53 (NTD_1-53_-sfGFP) residues of CsoSCA-NTD was co-expressed with synthetic carboxysomes in *E. coli* and purified to assess GFP encapsulation. Both NTD_1-37_-sfGFP and NTD_1-53_-sfGFP produced green fluorescent carboxysomes containing the fusion protein (Fig. 1G), while a negative control did not. Normalized for expression levels, the encapsulation efficiencies (sfGFP fluorescence of carboxysome/lysate) were as follows: NTD_1-37_-sfGFP: ∼3%, NTD_1-53_-sfGFP: ∼15% and CsoSCA-sfGFP: ∼23% (SI Table 2). This indicates that additional sequence elements not present in NTD_1-37_-sfGFP may be needed for efficient encapsulation. Shell and Rubisco protein levels were the same for all carboxysome constructs, and hence, should not influence the efficiency of NTD encapsulation (SI Table 2). In summary, these results demonstrate that the N-terminus is necessary and sufficient for encapsulating CsoSCA into the carboxysome.

### CsoSCA interacts with Rubisco

Previous immunogold labeling EM and biochemical assays (freeze/thaw treatment of carboxysomes) have suggested that CsoSCA may associate with the shell but no specific interactions have been described (20, 21). Thus, to identify CsoSCA’s interaction partner, we measured binding of un-tagged CsoSCA against all carboxysome proteins (the shell proteins CsoS1A, CsoS1B, CsoS1D, CsoS4B; the scaffolding protein CsoS2B; and Rubisco) using BLI. This screen showed that CsoSCA interacted with Rubisco, while none of the other carboxysome proteins had detectable binding above background (Fig. 2A). An N-terminal truncated protein variant (ΔNTD_1-37_CsoSCA) did not bind Rubisco, confirming NTD’s involvement in the interaction (SI Fig. 2A). Native-PAGE confirmed binding between CsoSCA and Rubisco and lack of binding to the major shell proteins CsoS1A and CsoS1B (SI Fig. 2B). Finally, co-elution of Rubisco and NTD_1-53_-sfGFP using size exclusion chromatography confirmed the interaction in a solution based assay (SI Fig. 2C).

**Figure 2:**
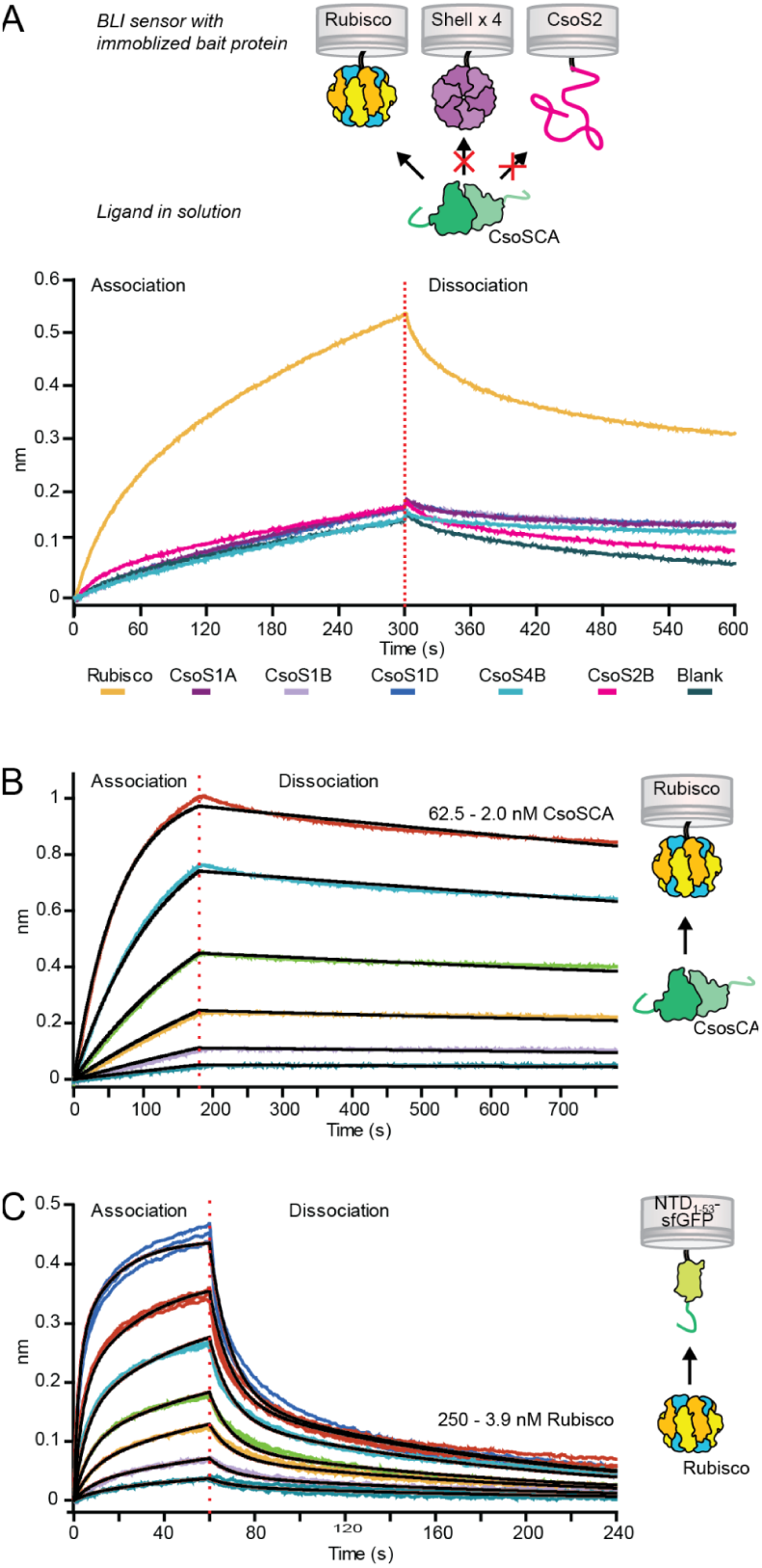
CsoSCA interacts with Rubisco via its N-terminal peptide. (A) Biolayer interferometry binding screen with CsoSCA against carboxysome proteins. Binding of CsoSCA (green) was assayed against Rubisco (yellow), the shell proteins; CsoS1A (purple), CsoS1B (light purple), CsoS1D (blue) and CsoS2B (light blue) and against the scaffolding protein CsoS2B (magenta). BLI responses showed binding to Rubisco, while none of the other carboxysome proteins showed detectable binding. (B) BLI response from binding affinity measurement of CsoSCA-MBP against immobilized Rubisco. CsoSCA concentration ranged from 62.5 - 2.0 nM in a 1:2 dilution series. (C) BLI response from binding affinity measurements of Rubisco against immobilized NTD_1-53_-sfGFP (light green). Rubisco concentration ranged from 250 - 3.9 nM in a 1:2 dilution series.

Concentration dependence of CsoSCA binding to Rubisco was confirmed by BLI and *K*_D_ of the interaction determined to be 1.2 nM ± 0.1 (*k*_on_= 2.5 x 10^5^ M^-1^s^-1^, *k*_off_= 2.9 x 10^-4^ s^-1^) (Fig. 2B and Table 1). Due to instability of wtCsoSCA, the *K*_D_ was measured with a CsoSCA-MBP fusion (the negative control of MBP alone did not bind Rubisco (SI Fig. 2D)). Although CsoSCA crystallized as a dimer (Fig. 1B) (22), our CsoSCA-MBP protein eluted on size exclusion chromatography at an estimated molecular weight of 612 kDa, consistent with a possible hexameric state (SI Fig. 2E,F). Rubisco binding to NTD_1-53_-sfGFP further confirmed the NTD_1-53_-peptide interaction (*K*_D1_ = 30 nM, *K*_D2_ = 80 nM; fit to a 1:2 model) (Fig. 2C, Table 1 and SI Fig. 2G). The 25-fold higher *K*_D_ with CsoSCA compared to NTD_1-53_-sfGFP is mainly an effect of a slower off-rate, demonstrating the importance of the multivalency (resulting from CsoSCA being multivalent while NTD_1-53_-sfGFP is monomeric) in obtaining a high affinity interaction with Rubisco.

### Structure of Rubisco in complex with CsoSCA NTD peptide

An emerging theme of CCM self-assembly is that Rubisco interacts with various CCM proteins via Short Linear Motifs (SLiMs) found in intrinsically disordered proteins/regions (IDP/Rs) (29–31). Since CsoSCA’s NTD appears to bind Rubisco (Fig. 2C and SI Fig. 3), we next sought to determine the structure of Rubisco in complex with this peptide using cryo-EM. In order to promote high occupancy of available binding sites, an excess of NTD peptide was complexed with Rubisco and imaged as described in the Materials and Methods. These data yielded two slightly distinct single particle reconstructions of Rubisco bound to a peptide corresponding to the first 50 residues of CsoSCA (NTD_1-50_) at 1.98 Å (State-1) and 2.07 Å (State-2) nominal resolution (SI Fig. 4 and 5, SI Table 3). Of note, both reconstructions show densities for ordered waters and alternate side-chain conformers and are of the highest resolution obtained by 200 keV cryo-EM to date (SI Fig. 5) (32). The two confirmations are highly similar (RMSD: 0.22 Å) with the most notable differences occurring in the β-sheet of the N-terminal domain of CbbL, in particular the conformations of loops P37-D42 and G115-G125 (SI Fig. 6). Since both reconstructions display similar density for the NTD_1-50_ peptide, we do not attribute the difference in confirmation to the presence of the peptide but rather subtle “breathing” of Rubisco. For clarity, we choose to predominantly focus our discussion on the higher resolution 1.98 Å structure (State-1).

**Figure 3:**
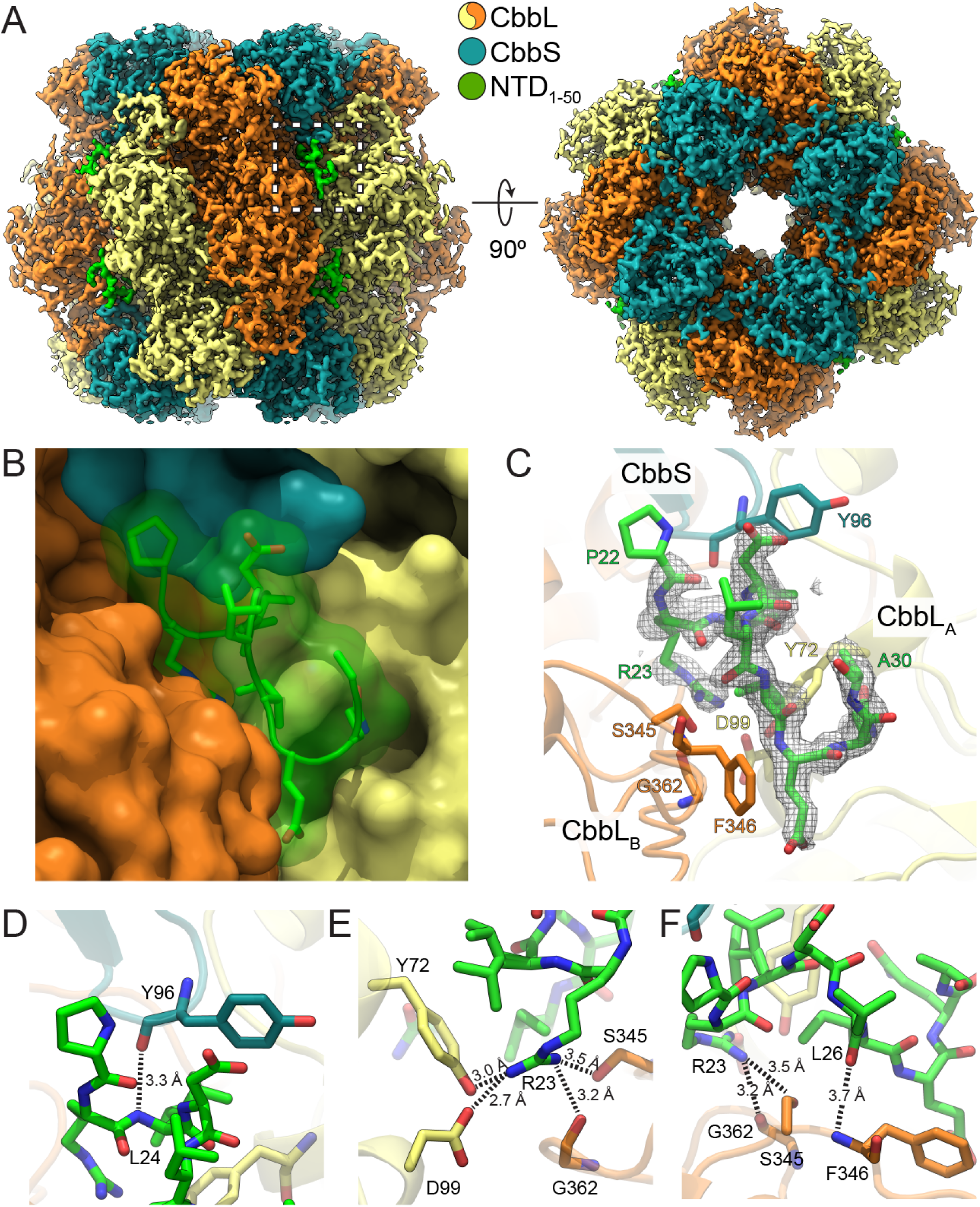
Structure of Rubisco with bound NTD_1-50_ CsoSCA peptide. (A) Cryo-EM map of Rubisco bound to a peptide corresponding to the first 50 residues of CsoSCA (NTD_1-50_). Rubisco-NTD_1-50_ co-complex is colored by subunit with color key inset. (B) Close-up of the region boxed in A of the NTD_1-50_ peptide shown as sticks and transparent surface and Rubisco subunits shown as opaque surfaces. (C) Same view as in B with NTD_1-50_ peptide and interacting Rubisco residues shown as sticks. NTD_1-50_ peptide density is shown as a grey mesh contoured to 2σ. (D-F) Detailed polar interactions between residues of NTD_1-50_ peptide and Rubisco are shown as sticks with interaction depicted as dashed lines with distances in Ångströms.

**Figure 4:**
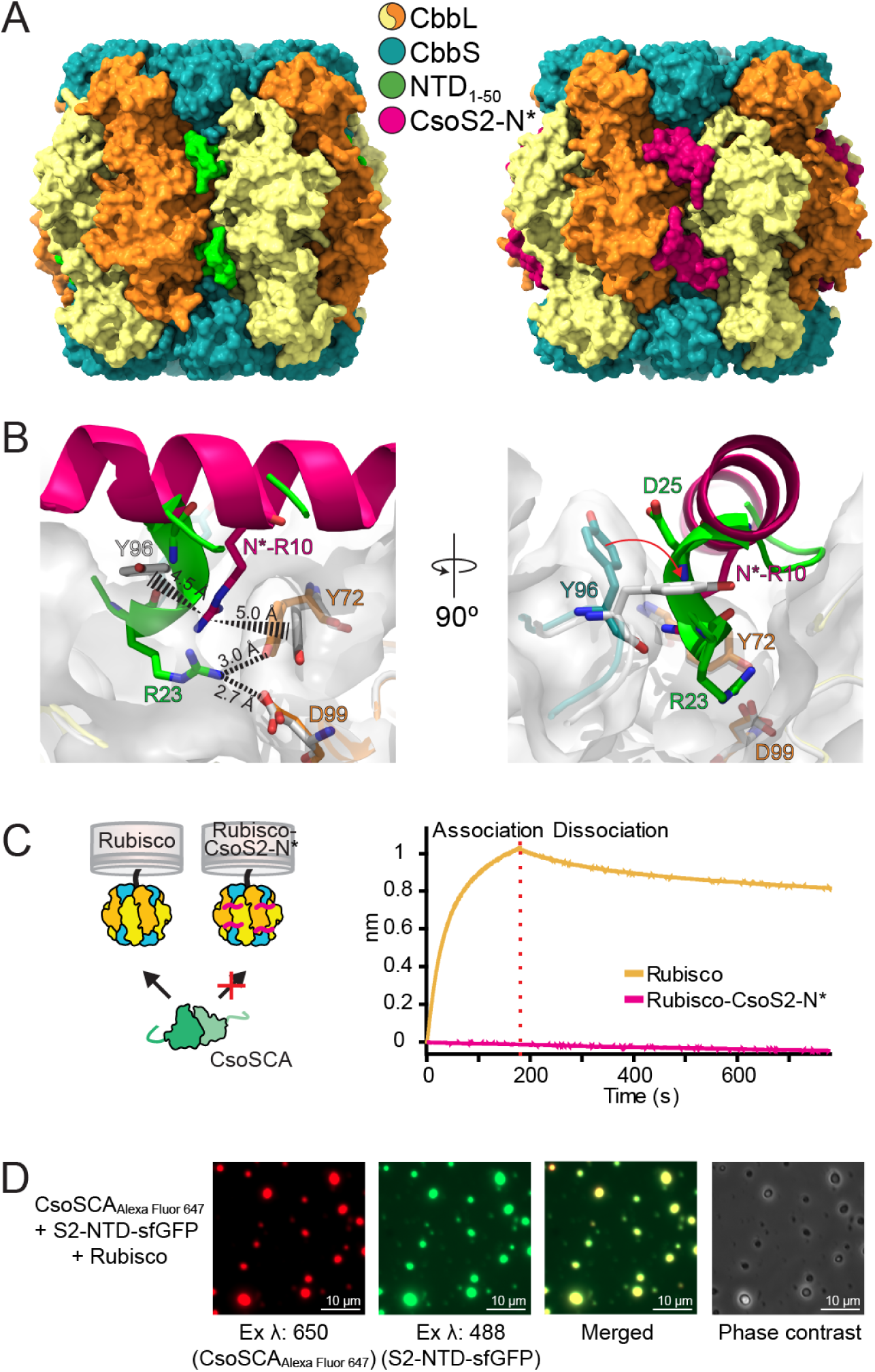
*CsoSCA and CsoS2* bind at the same site on Rubisco. (A) Surface representation of Rubisco with CsoSCA’s NTD_1-50_ peptide bound and Rubisco with CsoS2 peptide bound (pdb: 6uew). (B) Zoomed view of binding site showing the different conformations of the CsoSCA and CsoS2 peptides. Peptides are shown as cartoons and detailed residues as sticks. Rubisco bound with the NTD_1-50_ peptide is colored according to color key inset in (A) and Rubisco (both subunits) bound with CsoS2-N* peptide (pdb: 6uew) is colored white. White transparent surface represents the Rubisco structure which binds CsoSCA. Polar interactions are depicted as dashed lines and cation-pi stacking as dashed triangles with distances in Ångströms. (C) BLI response shows that the CsoS2 peptide fused to Rubisco (Rubisco-CsoS2-N*) passivates binding of CsoSCA to Rubisco. (D) Alexa Fluor 647, sfGFP and merged fluorescence as well as phase contrast images of protein condensates formed from a solution of Rubisco, CsoS2-NTD-sfGFP and Alexa Fluor 647 labeled CsoSCA-MBP showing that CsoSCA recruits into Rubisco-CsoS2 protein condensates.

**Figure 5:**
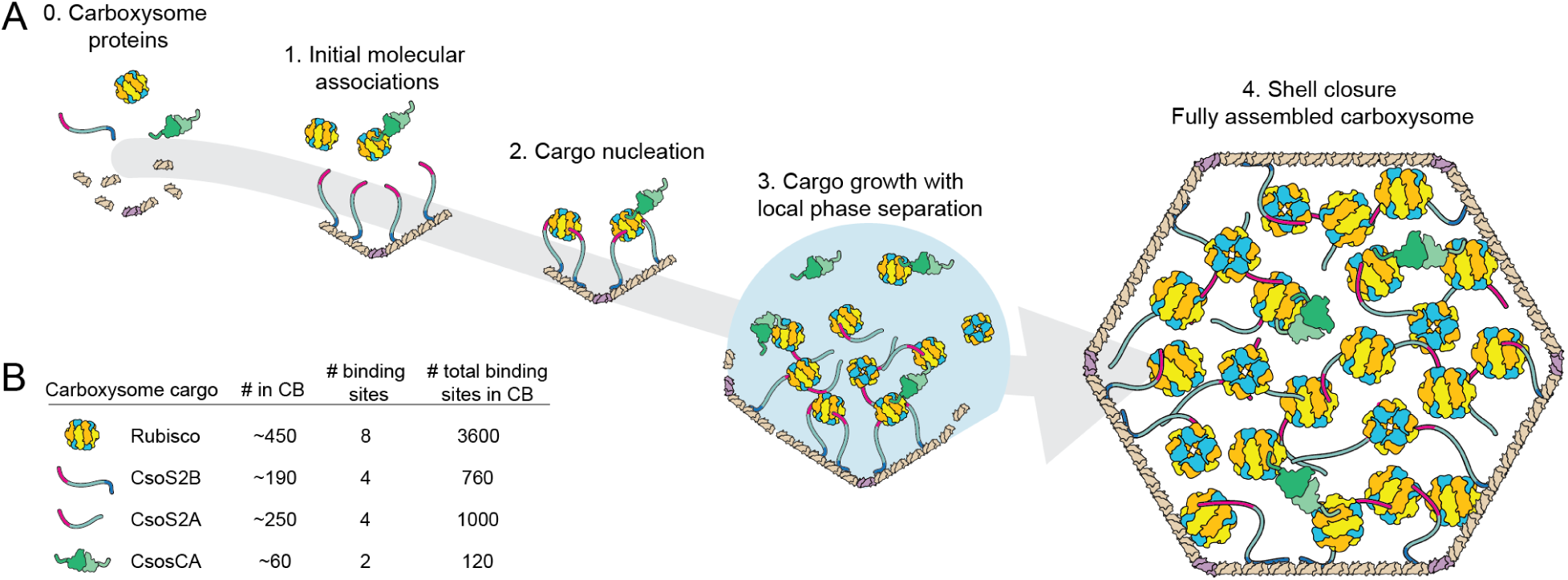
Updated model for carboxysome assembly. (A) Schematic model of α-carboxysome assembly in which CsoSCA is recruited to the carboxysome via interactions with Rubisco. The model involves 1. Initial molecular associations (specific order not known), 2. Cargo nucleation, 3. Cargo growth with local phase separation and finally 4. Shell closure forming fully assembled carboxysomes. Current knowledge does not allow us to distinguish between whether CsoSCA associates with Rubisco during the initial association, step 1, or if association occurs in the phase separated condensate, step 3 (or both). The fully assembled carboxysome in step 4 shows a stoichiometrically accurate - with respect to cargo proteins - version of the α-carboxysome. (B) Average number of cargo proteins present in an α-carboxysome, number of binding sites per oligomeric form of cargo protein, and total number of binding sites per carboxysome (35).

In the cryo-EM reconstruction of the Rubisco-NTD_1-50_ complex, density corresponding to NTD_1-50_ is located in a groove formed at the interfaces of two CbbL subunits (from two different CbbL_2_ dimers) and one CbbS (Fig. 3). The biological assembly of Rubisco is CbbL_8_S_8_, resulting in eight of these binding sites per Rubisco oligomer. The peptide density is of marginally lower quality (local resolution estimate: 2.1-2.2 Å) than that of the surrounding Rubisco density. Nevertheless, we could confidently assign the density to nine residues of the NTD_1-50_ peptide starting at P22 (PRLDLIEQA) (Fig. 3B,C). The structure of the Rubisco-NTD_1-50_ complex is highly similar to a previous crystal structure of Rubisco (PDB: 1SVD, RMSD: 0.45 Å, SI Fig. 7), indicating that binding of the NTD_1-50_ peptide does not induce large-scale conformational changes in Rubisco.

The resolved region of the peptide starts at the bottom of the CbbS subunit (around loop D94-S99), runs downward within the groove between the two CbbL subunits and ends between β-strand S345-I347 in CbbL_B_ and loop P19-I29 in CbbL_A_, that results in a buried interface of approximately 700 Å^2^. This short stretch of sequence is predicted by JPred to form an alpha-helix (SI Fig. 8A). Indeed, we observe this segment to form a single helical turn and thereafter an extended coil (Fig. 3B,C). The sharp turn of the backbone introduced by P22 (the first observed residue of the bound peptide) ensures that the upstream peptide chain points outwards towards the solvent instead of clashing with CbbL.

### Binding is predominantly mediated by a network of hydrogen bonds

Our atomic model of the Rubisco-NTD_1-50_ co-complex indicates that the interaction between the NTD_1-50_ peptide and Rubisco is largely mediated through polar interactions and, predominantly, hydrogen bonds. R23 forms an extensive network of interactions with the neighboring CbbL subunits. The side chain of R23 forms a salt-bridge with D99 (CbbL_A_) and hydrogen bonds to the hydroxyl groups of the CbbL subunits Y72 (CbbL_A_) and S345 (CbbL_B_), as well as to the carbonyl of G362 (CbbL_B_) (Fig. 3E). The L24 amide and L26 carbonyl hydrogen bond to the carbonyl of CbbS Y96 and amide of CbbL_B_ F346, respectively (Fig. 3D,F).

A water-mediated hydrogen bonding network likely also contributes to peptide binding (SI Fig. 9). This putative network is predominantly built up by backbone-water interactions and consists of interactions between N29 and CbbL_A_ Y72 (SI Fig. 9A,B), R23 and CbbL_B_ S345, and L26 and CbbL_B_ F346 (SI Fig. 9C). However, in the lower resolution structure (State-2) the two waters mediating the interactions between the peptide and CbbL_B_ are not resolved (SI Fig. 9D), possibly due to the slightly lower resolution of this reconstruction. Rubisco residues interacting with CsoSCA have a high conservation score among α-carboxysomal Rubiscos, but are in general not conserved in β-carboxysomal Rubiscos (SI Fig. 8B).

To determine the relative importance of the different interactions, we measured binding kinetics with a selected set of point mutations on both the NTD peptide and Rubisco (Table 1, SI Table 4 and SI Fig. 10). The P22A mutation resulted in a dramatic loss in binding. While a protonated CbbS D94 could potentially hydrogen bond with the amide of P22, this large effect is more likely due to P22’s importance in establishing the initial alpha-helical backbone conformation of the peptide or the sharp backbone turn that is essential for binding.

Despite the many interactions made by the buried peptide residue R23, mutation of this residue to alanine yielded roughly the same *K*_D_-value as wild-type. However, mutation of the residues on CbbL_A_ which interact with R23 - Y72A and D99A - resulted in a 10-fold increase in *K*_D_ (mainly an effect of slower on-rate). These results are consistent with the net contribution of interactions made by R23 to binding to be quite low, but, nevertheless, this residue adversely affects binding when these interactions are not satisfied in a buried conformation. In this context, R23 may play a role in establishing the specificity of the interaction between Rubisco and CsoSCA.

The remaining hydrogen bonds between the NTD peptide and Rubisco are mediated by backbone moieties. Furthermore, the peptide is tightly-packed in the cleft formed by Rubisco to form a buried interface comprising 700 Å^2^ of the 1200 Å^2^ solvent accessible surface area of the peptide. Thus, given the strong negative effect of the P22A mutant on binding, shape-complementarity between peptide and Rubisco appears critical to enable the extensive backbone-mediated hydrogen bonds and van der Waals interactions that drive binding.

### CsoSCA binds at the same site as CsoS2

α-carboxysome assembly is mediated by a repetitive and disordered protein, CsoS2, which is thought to bind both Rubisco and shell proteins, thus serving as a physical scaffold bridging these two major components. We previously solved the structure of Rubisco in complex with an N-terminal peptide derived from CsoS2 (CsoS2-N*) (29). Surprisingly, CsoSCA and CsoS2 bind at nearly the same location on Rubisco but utilize substantially different SLiMs and binding modes (Fig. 4A,B).

CsoS2-N* is largely alpha-helical and binds Rubisco by spanning over the CbbL_2_ dimer interface lying on top of the protein surface (Fig. 4B). The complex is highly dependent on salt bridges and cation-pi interactions. In contrast, the CsoSCA peptide is bound in a conformation turned roughly ∼45 degrees and with a greater fractional buried surface area for the observed peptide (approximately 700 Å^2^ of 1200 Å^2^ solvent accessible surface area, compared to 830 Å^2^ of 2500 Å^2^). CsoSCA is buried deeper into the groove between the two CbbL subunits and interacts mainly via hydrogen bonds and what appears to be an ordered network of water molecules. Notably, both peptides make significant interactions with Rubisco CbbL Y72 (Fig. 4B). This residue is conserved in α-carboxysome Rubisco, but not in Rubisco from β-carboxysomes or the Form II Rubisco in *H. neapolitanus,* and likely contributes to specificity. Both proteins interact with Y72 via arginines; however, in CsoS2-N* R10 is cation-pi stacked between CbbL Y72 and CbbS Y96, while in CsoSCA, R23 is positioned deeper into the structure and hydrogen bonds with CbbL Y72 and D99. Another notable feature is CbbS Y96, which in the Rubisco-CsoS2 structure is flipped ∼90 degrees compared to the wild-type and Rubisco-CsoSCA structure (Fig. 4B), covering the groove between the CbbL subunits interface and enabling the conformation necessary for cation-pi interaction.

Combined, these interactions would seemingly make it impossible for CsoSCA and CsoS2 to bind to the same site of Rubisco at the same time. We have previously developed a Rubisco-CsoS2-N* fusion with all such binding sites occupied due to high local concentration of the CsoS2-N* peptide. As expected, BLI measurement indicated that NTD_1-53_-sfGFP cannot bind to Rubisco when CsoS2-N* is already present, thus confirming that CsoS2 and CsoSCA compete for the same binding site (Fig. 4C). Earlier experiments from our group have demonstrated *in vitro* condensate formation between Rubisco and CsoS2-NTD suggesting that assembly of the ɑ-carboxysome occurs through a condensation-like event (29). Here, we extended these experiments to include CsoSCA in the Rubisco-CsoS2-NTD condensates. These results clearly show that CsoSCA is recruited into the phase-separated Rubisco-CsoS2-NTD condensates (Fig. 4D) and demonstrate that all three proteins can simultaneously participate in such a protein interaction network.

## Discussion

In this study, we have determined the structural basis for carbonic anhydrase encapsulation in α-carboxysomes. We found that in the model organism *H. neapolitanus*, CsoSCA and Rubisco form a previously unknown supercomplex. Through biophysical measurements and *in vivo* experiments we found that this complex-formation is dependent on the intrinsically disordered N-terminal peptide of CsoSCA. The cryo-EM structure of Rubisco in complex with this peptide reveals that CsoSCA binds Rubisco at a site overlapping with that of the scaffolding protein CsoS2. Aside from its enzymatic activity, this establishes Rubisco’s additional function as an interaction hub in the assembly of the α-carboxysome.

### Updated model of α-carboxysome architecture and assembly

The intrinsically disordered and repetitive protein CsoS2 acts as a scaffold between shell and Rubisco and orchestrates the assembly (33, 34) of the α-carboxysome. We previously discovered that a repeat of conserved SLiMs (four repeats/protein) found in the N-terminal portion of CsoS2 binds Rubisco and is essential for carboxysome formation (29). Here we demonstrate that the carboxysomal carbonic anhydrase (CsoSCA) is recruited to the carboxysome via association with Rubisco. We further show that CsoSCA’s N-terminal targeting peptide and CsoS2 bind at the same site on Rubisco. Figure 5A presents our current model of α-carboxysome assembly.

Previous work has shown that, on average, there are 450, 440 and 60 copies of Rubisco, CsoS2 and CsoSCA in a typical *H. neapolitanus* carboxysome, respectively (Fig. 5B) (35). The roughly 1800 CsoS2 and 120 CsoSCA Rubisco binding motifs per carboxysome set an upper boundary of occupancy for the binding sites of Rubisco (∼3600 sites/carboxysome). Assuming all CsoS2 and CsoSCA motifs engage in binding, roughly 50% of Rubisco sites would be occupied by CsoS2 while considerably less, ∼3.5%, would be occupied by CsoSCA. Although this assumes that all Rubisco sites are accessible and that all CsoS2 and CsoSCA motifs bind, both assumptions of which could be incorrect, such a calculation indicates there is likely a surplus of Rubisco sites available for binding. Our finding that CsoSCA is recruited to Rubisco-CsoS2-NTD condensates supports the hypothesis that this ternary complex is an important feature of cargo assembly in ɑ-carboxysomes *in vivo* (Fig. 4D and SI Fig. 11). Further, it has previously been shown that CsoSCA mRNA levels are lower compared to other carboxysome genes (36), suggesting that the amount of encapsulated CsoSCA is likely regulated by protein expression level rather than by competing for binding site occupancy with CsoS2. Due to the need for tight regulation of CA activity outside of the carboxysome, efficient encapsulation is vital (19) and a scenario where CsoSCA had to compete for binding could pose a physiological problem.

One specific unknown is the importance of CsoSCA multivalency imparted by its oligomeric structure and the resulting mode of interaction with Rubisco. The significantly slower dissociation rate of full-length CsoSCA (multivalent) compared to the NTD_1-53_-peptide (monovalent) implies an importance of multivalent protein-protein interactions (Table 1 and Fig. 2B,C), a feature commonly observed for other IDP/R involved in phase separation (37). In terms of binding mode, bivalent binding of full-length CsoSCA could occur either between two binding sites on the same Rubisco or between two sites on different Rubisco molecules. The relatively short stretch of IDR sequence before the Rubisco binding motif and the rigidity of the folded domains presumably constrains possible binding conformations where CsoSCA binds on top or on the side in a 1:1 CsoSCA-Rubisco complex (SI Fig. 12). Alternatively, CsoSCA could cross-link two Rubisco molecules (SI Fig. 12). Further, previous experiments have indicated that CsoSCA is localized to the shell (20, 21). Although our data suggest that CsoSCA makes a primary interaction with Rubisco, and would likely be found throughout the carboxysome, it is possible that additional unknown protein interactions could bias CsoSCA localization towards the shell. Recent cryo-electron tomography work has been unable to unambiguously locate CsoSCA inside the carboxysome (38, 39). However, rapid advances in this technique will likely, in the near future, determine the binding conformation and localization of all such components *in situ*.

### Plasticity of the Rubisco binding motif

We could not identify a consensus Rubisco SLiM binding motif across NTD sequences in CsoSCA homologs. The binding element identified in *H. neapolitanus* CsoSCA (PRLDLIEQA) is present in its most closely related homolog (*Halothiobacillus sp. LS2*) but is not conserved across species (further discussed in SI Fig. 13A,B). Many Prolines are followed by R or xR, but, overall, the proteobacteria clade contains no convincing conserved motifs. In the cyanobacterial clade, a PTAPx[R/K]R motif is present in 87% of the sequences suggesting a possible binding motif among cyanobacteria.

Surprisingly, a handful of cyanobacterial CsoSCA sequences from the *Procloroccocos* genus contain the Rubisco-binding motif found in CsoS2 (RxxxxxRRxxxxxxGK) (SI Fig. 13A), suggesting an evolutionary relationship. The lack of a consensus motif in CsoSCA homologs, coupled with the fact that CsoSCA and CsoS2 bind at the same site but with different mechanisms, reveals an evolutionary plasticity in SLiM sequence space. Across the various microbes in this phylogeny, we hypothesize that CA is recruited to its respective α-carboxysomes by the observed, versatile Rubisco binding site and does so using diverse SLiM sequences.

### Rubisco as an interaction hub in biophysical CCM’s

Recent work on both bacterial carboxysomes and algal pyrenoids suggests that Rubisco itself plays a role as an interaction hub in the ultrastructural organization of CCMs. It is now clear that not only does Rubisco interact with scaffolding proteins as a means to condensate Rubisco and form these confined CO_2_-fixing organelles, it also recruits auxiliary proteins, such as CAs and activases, needed for the CCM to function. In the bacterial carboxysomal α-lineage, we have demonstrated that Rubisco binds the intrinsically disordered proteins CsoS2 (29) and CsoSCA. Additionally, it also binds the Rubisco activase CbbQO, likely via CbbO’s von Willebrand factor A domain (40, 41). The β-carboxysomal Rubisco binds its interaction partners - the scaffolding protein CcmM (42, 43), and the Rubisco activase Rca (44, 45) - via a folded domain resembling the small subunit of Rubisco (SSLD). Similar to CsoSCA, the β-carboxysomal CA, CcaA, is also recruited via a terminal peptide. However, instead of direct interaction with Rubisco, the two enzymes are linked together via the scaffolding protein CcmM (28). This convergent function may have evolutionary significance - recent results suggest that Rubisco-CA co-localization was an important step in the evolution of biophysical CCMs (46, 47). Despite convergent evolution, a notable similarity between both carboxysome lineages is the binding site on Rubisco. In known cases (except for CbbQO), the interactor binds at the same patch on Rubisco and makes contact with two different CbbL_2_ dimers and one CbbS. This likely ensures that the binding partner only interacts with fully assembled CbbL_8_S_8_ Rubiscos during the assembly process.

In contrast to these bacterial systems, in the model algae *Chlamydomonas reinhardtii* a repeat SLiM in the disordered scaffolding protein EPYC1 is essential for pyrenoid formation (48, 49) and binds on top of the small subunit via salt bridges and a hydrophobic interface (30). The sequence motif is shared among many pyrenoid proteins, suggesting a mechanism for protein targeting as well as more broadly organizing pyrenoid ultrastructure (31). The versatility in binding motif and binding site, and the convergent function of diverse Rubiscos as a hub of interaction implies this might be a general feature, which raises a final question: do Form IB plant Rubiscos engage in similar protein-protein interactions and do other Rubisco Forms also function as interaction hubs?

In summary, this work advances our understanding of carboxysome biogenesis, puts a focus on the essential carbonic anhydrase, and the central role of Rubisco as a hub protein. This provides critical findings for engineering the carboxysome based CCM into *e.g.* crops and industrially relevant microorganisms for improved growth and yields. More broadly, we hope that the findings presented here will advance our understanding of bacterial microcompartments and promote development of their many potential biotechnological applications.

## Materials and Methods

### Bioinformatics

Protein sequences assigned CsoSCA (pfam08936) from all finished and permanent draft bacterial genomes available in the Integrated Microbial Genomes and Microbiomes database (IMG) (50) were collected on December 12, 2019 and curated to only include proteins in an α-carboxysome operon (containing CbbL/S, CsoS2 and shell hexamers and pentamers) (412 genes). Thereafter redundancy was reduced by removing sequences with >98% identity using Jalview and sequences were manually curated to remove incomplete sequences, resulting in 222 sequences in the final CsoSCA dataset (SI Table 1). Sequences were aligned using MUSCLE (51). Resulting MSA was used to calculate conservation score (52). Tree was built using IQ-TREE web server (53) and visualized using iTOL (54). Protein disorder were predicted for a subset of the dataset, including *H. neapolitanus* CsoSCA, using the DISOPRED3 algorithm (55). Conservation of CsoSCA NTD Rubisco binding motif was analyzed using The MEME Suite (56) (SI Table 1). MSA of α-carboxysomal Form IA Rubiscos (135 cbbL and 132 cbbS sequences) and β-carboxysomal Form IB Rubiscos (211 cbbL and 207 cbbS sequences) was constructed (SI Table 1) using, MUSCLE and visualized with WebLogo. Secondary structure prediction of NTD sequence was performed using Jpred4 (57).

### Protein expression and purification

Specific details regarding *E. coli* strain, plasmid, and expression conditions for each protein in this study (CsoSCA variants, sfGFP-fusions, Rubisco, shell proteins and CsoS2) are provided in SI Table 5. Expression plasmids were transformed into either *E. coli* BL21-A1 or BW25113 cells (SI Table 5). For protein expression, cells harboring appropriate plasmids (SI Table 5) were grown at 37 °C in LB-medium supplemented with appropriate antibiotics. All Rubisco constructs were co-transformed with pGro7 for co-expression of GroEL/ES. At OD_600_ = 0.4 - 0.6 the temperature was decreased to 18 °C and expression induced by addition of 0.08% L-arabinose, 0.1 μM anhydrotetracycline (ATc) or 0.5 mM IPTG, see SI Table 5. Cells were grown overnight, harvested by centrifugation at 5,000 x *g* and frozen at -20 °C until use. All proteins were purified with either His-tag och Strep-tag affinity purifications. Bacterial pellets were thawed and resuspended in appropriate binding buffer supplemented with 0.2 mM phenylmethanesulfonyl fluoride (PMSF), 0.1 mg/mL lysozyme and 0.1 μL/mL benzonase and lysed by three passes through a homogeniser (Avestin EmulsiFlex-C3) or by chemical lysis by diluting the resuspension 1:1 in B-PER II (Thermo Fisher) an incubating in RT for 30 min on a rocking table. Lysed cells were clarified by centrifugation at 27,000 x *g* for 45 min.

A summary of purifications of the proteins used in this study is described below. Specific details are provided in SI Table 5. *His-tag purification (used for all CsoSCA-variants, sfGFP-fusions, shell proteins and CsoS2, see SI Table 5):* Clarified lysate was applied to a 5 mL HisTrap FF column (GE Healthcare) equilibrated with His-binding buffer (50 mM Tris, 300 mM NaCl, 20 mM imidazole, pH 8.0) using a syringe pump at a flow rate of 5 mL/min. Unspecific bound proteins were washed away with His-buffer containing 60 mM imidazole until A280 reached a stable baseline, and protein thereafter eluted with His-buffer containing 300 mM imidazole. *Strep-tag purification (used for Rubisco-variants, see SI Table 5):* Clarified lysate was applied to a 5 mL StrepTrap column (GE Healthcare) equilibrated with Strep-buffer (50 mM Tris, 300 mM NaCl, pH 8.0). Unspecific bound proteins were washed away with Strep-buffer until A280 reached a stable background, and protein thereafter eluted with Strep-buffer supplemented with 2.5 mM *d*-desthiobiotin (Sigma-Aldrich). All purified proteins were concentrated to appropriate concentration using 30 kDa Mw cut-off centrifugal filters (Amicon) and buffer exchanged into 50 mM Tris, 150 mM NaCl, pH 7.5 using 10DG Desalting Columns (Bio-Rad). *Untagged CsoSCA:* Untagged CsoSCA were purified as a His-SUMO fusion as outlined above. After elution from HisTrap column His-SUMO-CsoSCA was concentrated from 10 mL to 3 mL and buffer exchange into 50 mM Tris, 300 mM NaCl, 10% glycerol pH 8.0. The SUMO-tag was thereafter cleaved by addition of 1:200 molar ratio of Ulp-protease:His-SUMO-CsoSCA and 1 hour incubation. Cleaved protein mix was applied to a 1 mL HiTrap column to remove His-SUMO and residual uncleaved His-SUMO-CsoSCA. This resulted in a pure and untagged CsoSCA. Since full-length CsoSCA is prone to crash out in high protein concentrations care was taken to not exceed CsoSCA protein concentration above 1 mg/mL. *CsoSCA-MBP:* After His-tag purification protein was subsequently purified by size exclusion chromatography using a Superose 6 Increase 10/300 column (GE Healthcare). The oligomeric state of CsoSCA-MBP was determined from the Superose 6 Increase chromatogram and a calibration standard (Bio-Rad). Molecular weight of standard was as follows: 1. Thyroglobin (670 kDa) 2. γ-globulin (158 kDa) 3. Ovalbumin (44 kDa) 3. Myoglobin (17 kDa) Vitamin B12 (1.4 kDa). Protein purities were assessed by SDS-page and were in general >95% pure. Protein concentrations were determined by A_280_ and the theoretically calculated extinction coefficient (ProtParam). All purifications were carried out at 4 °C. For storage, proteins were made to 10% (w/v) glycerol, flash-frozen in liquid nitrogen and stored in -80 °C.

### Growth phenotypes of *H. neapolitanus csoSCA* mutants

#### Generation of H. neapolitanus ΔcsoSCA and of WTcsoSCA and ΔNTD_1-49_csoSCA mutant complementations

*csoSCA* was knocked out by transforming a plasmid containing a spectinomycin cassette with 1KB homology arms adjacent to *csoSCA* gene. Colonies that grew on spectinomycin were verified by colony PCR and Sanger Sequencing to verify a clean knockout. CsoSCA mutant complementations (*ΔcsoSCA+WTcsoSCA; ΔcsoSCA+NTD_1-49_csoSCA)* were generated by transformation and concomitant genomic integration via homologous recombination into *H. neapolitanus* NS2 neutral site and verified by colony PCR and Sanger sequencing.

#### H. nea growth assays

Pre-cultures of WT *H. neapolitanus* and *H. neapolitanus ΔcsoSCA* were grown in DSMZ68 at 5% CO_2_. *ΔcsoSCA* transformed with wild-type *csoSCA* or the N-terminal truncation *ΔNTD_1-49_CsoSCA* were cultured in the same conditions with the addition of 1 μM IPTG to induce CA expression. All precultures were grown with the appropriate antibiotics. Upon reaching log phase, cultures were spun down for 15 minutes at 4,000 *x g*. Pellets were then resuspended in 1 mL DSMZ68 without thiosulfate and pH indicator. Cultures were transferred into 1.7 mL eppendorf tubes and centrifuged at 4,000 x *g* for 8 minutes. This wash step was repeated twice before the cultures were diluted 5x and then normalized to a cell density of 0.1 OD600. Normalized cultures were serially diluted in 10x steps from 10^-1^ to 10^-8^ OD600. Resulting titers were spotted in 3 μL volumes onto plates with appropriate antibiotics in 5% CO_2_ and ambient air; strains expressing complemented *WTcsoSCA* or *ΔNTD_1-53_csoSCA* were plated on plates containing 1 μM IPTG. Strains were allowed to grow for 4 days. All strains were plated in biological and technical triplicate.

### Carboxysome purifications

#### Small scale enrichment of carboxysomes

100 mL of *E. coli* BW25113 cells harboring pHnCB10 (plasmid for homologous expression of *H. neapolitanus* α-carboxysomes in *E. coli*), containing either wildtype, ΔNTD_1-37_ or ΔNTD_1-49_ truncation of flag-tagged CsoSCA, were grown at 30 °C in LB-medium supplemented with appropriate antibiotics (SI Table 5). At OD_600_ = 0.4 - 0.6 the expression was induced by addition of 0.5 mM IPTG, cells grown for 4 h, harvested by centrifugation at 5,000 x *g* and frozen at -20 °C until use. To enrich carboxysomes, cells from 100 mL of culture were resuspended and chemically lysed for 30 min in 6 mL of B-PER II (Thermo Fisher) diluted to 1x with TEMB buffer (10 mM Tris pH 8.0, 10 mM MgCl2, 20 mM NaHCO_3_ and 1 mM EDTA) supplemented with 0.1 mg/mL lysozyme, 1 mM PMSF and 0.1 μL of benzonase/mL (Sigma-Aldrich). Lysed cells were centrifuged 12,000 x *g* for 15 min to remove cell debris. The clarified lysate was centrifuged 40,000 x *g* for 30 min and the enriched carboxysome fraction were thereafter resuspended in 200 μL of TEMB.

#### Full scale purification of carboxysomes co-expressed with CsoSCA-sfGFP fusions

1 L of *E. coli* BW25113 cells co-transformed with pHnCB9 (pHnCB10 lacking the gene for *csoSCA*) and pFA-plasmid containing a CsoSCA variants fused to sfGFP (NTD_1-37_-sfGFP, NTD_1-53_-sfGFP or CsoSCA-sfGFP) were grown at 37 °C in LB-medium supplemented with appropriate antibiotics (SI Table 5). At OD_600_ = 0.4 - 0.6 the expression was induced by addition of 0.5 mM IPTG and 0.1 μM aTc, temperature decreased to 18 °C and grown o/n. Thereafter cells were harvested by centrifugation at 5,000 x *g* and frozen at -20 °C until use. To purify carboxysomes, cells were chemically lysed for 30 min under mild shaking in B-PER II (Thermo Fisher) diluted to 1x with TEMB buffer supplemented with 0.1 mg/mL lysozyme, 1 mM PMSF and 0.1 μL of benzonase/mL (Sigma-Aldrich). Lysed cells were centrifuged 12,000 x *g* for 15 min to remove cell debris. The clarified lysate was centrifuged 40,000 x *g* for 30 min to pellet carboxysomes and obtained pellets were gently resuspended in 1.5 mL TEMB buffer. Resuspended pellets were loaded on top of a 25 mL 10-50% sucrose step gradient (10, 20, 30, 40 and 50% w/v sucrose, made in TEMB buffer) and ultracentrifuged at 105,000 x *g* for 35 min (SW 32 Ti Swinging-bucket, Beckman Coulter). Gradients were fractionated, analyzed by SDS-PAGE and carboxysome containing fractions pooled and ultracentrifuged 100,000 x *g* for 90 min. Resulting pellets were gently resuspended in TEMB to obtain the final purified carboxysome sample.

Final carboxysome samples and as well as the lysate were analyzed for presence of CsoSCA or sfGFP-fusion protein by SDS-page (4–20% Mini-PROTEAN® TGX™ Precast Protein Gels (Bio-Rad)), western-blot and GFP-fluorescence. For western blotting, proteins from SDS-page gels were transferred to nitrocellulose membranes using the Trans-Blot Turbo system (Bio-Rad). Membranes were blocked with 5% (w/v) non-fat dry milk in phosphate-buffered saline (PBS), 0.1% (v/v) Triton X-100 for one hour at room temperature. Immunolabeling of Flag-tag was done overnight in 4 °C in the above mentioned buffer containing a 1:5,000 dilution of a monoclonal anti-Flag horseradish peroxidase conjugated antibody (Sigma). Membranes were washed 3×10 minutes with PBS, 0.1% (v/v) Triton X-100, and blots were thereafter developed using SuperSignal West Pico Chemiluminescent Substrate (ThermoFisher) according to manufacturer’s procedure. Gels and western blots were imaged with ChemiDoc^TM^ XRS+ System (Bio-Rad). Fluorescence of sfGFP-samples was quantified using a Infinite M-1000 plate reader (Tecan). To quantify efficiency of encapsulation between the different samples the ratio of sfGFP fluorescence in: carboxysomes (encapsulated protein)/lysate (expressed protein), was used. The ratio of shell/Rubisco content was quantified using densitometry by measuring the intensity of the CsoS1B and CbbS bands on the SDS-page using ImageJ.

### Biolayer interferometry

Protein-protein interactions were measured by Biolayer interferometry (BLI) using an Octet RED384 (Forte Bio). Experimental binding sequence used was: Loading bait 60-240 s, buffer wash 60 s, prey association and prey dissociation (followed by sensor regeneration for Ni-NTA sensors). *CsoSCA binding screen*: Purified Rubisco, CsoS1A, CsoS1B, CsoS1D, CsoS4B and CsoS2B were used as bait proteins and screened for CsoSCA binding (see purification protocol above). Bait proteins were immobilized on Octet® Ni-NTA Biosensors (Forte Bio) via terminal His-tag using: 5 μg/mL Rubisco, 3.5 μg/mL SUMO-S1A, 8 μg/mL SUMO-S1B, 5 μg/mL SUMO-S1D, 1.3 μg/mL SUMO-S4B or 2.6 μg/mL CsoS2B. To avoid tiling of shell proteins on the sensor surface SUMO-fusions were used for CsoS1A, 1B, 1D and 4B (58). 1 μM untagged CsoSCA was used as soluble prey protein. Binding assay was performed in a final buffer of 30 mM Tris pH 7.5, 145 mM NaCl, 1.0 mM TCEP and 0.01% Triton X100. After a run Ni-NTA sensors were regenerated in 50 mM Tris pH 8.0, 300 mM NaCl, 300 mM imidazole, 0.05% (w/v) SDS and experiment performed in triplicate. *NTD_1-53_-sfGFP vs. Rubisco:* Assays were performed as described above using 5 μg/mL NTD_1-53_-sfGFP-His as bait and varied concentration of strep-Rubisco as prey (wt: 250 - 3.9 nM, for mutants se Fig. SI 10) in 25 mM Tris pH 7.5, 70 mM NaCl and 0.01% Triton X100 and performed in triplicate of duplicates. When indicated, point mutants of NTD_1-53_-sfGFP or Rubisco were used. *Rubisco vs. CsoSCA*: 2.5 mg/mL of biotinylated Rubisco was immobilized as bait protein on Octet® Streptavidin Biosensors (Forte Bio) and 62.5 - 2.0 nM of C-terminal MBP tagged CsoSCA was used as prey. Experiment was performed in triplicate in 25 mM Tris pH 7.5, 125 mM NaCl and 0.01% Triton X100. *Rubisco vs. CsoSCA’s NTD_1-50_ peptide*: Biotinylated Rubisco was used as bait and the NTD_1-50_ peptide (100, 50 and 10 μL) as prey in a final buffer of 25 mM Tris, pH 7.5, 85 mM NaCl and 0.01% Triton X100. Binding and kinetic constants were extracted using the Data Analysis HT 10.0.00.44 software in the Octet Forte Bio package. NTD_1-53_-sfGFP vs. Rubisco were fitted to a 1:2 (Bivalent Analyte) binding model and Rubisco vs. CsoSCA to a 1:1 binding model.

### Size exclusion chromatography analysis of **NTD_1-53_**-sfGFP:Rubisco co-complex

Purified Rubisco-strep and NTD_1-53_-sfGFP samples from above were exchanged into moderate-salt buffer (20 mM Tris, pH 7.5, 150 mM NaCl) using Zeba desalting columns. For co-complexing, the protein samples were mixed at an 32:1 ratio (NTD_1-53_-sfGFP:CbbL_8_S_8_) and incubated briefly on ice prior to injection over a 3.2/300 Superose 6 Increase column equilibrated in 20 mM Tris, pH 7.5, 150 mM NaCl at 4 °C. The column was eluted isocratically in the same buffer with elution of total protein monitored by absorbance at 280 nm and elution of sfGFP-containing fractions monitored by absorbance at 485 nm.

### Native-page analysis of CsoSCA binding

Binding of CsoSCA to Rubisco and shell proteins was analyzed by native-page, using 4–15% Mini-PROTEAN® TGX™ Precast Protein Gels (Bio-Rad, USA). 2.5 μM CsoSCA was mixed with 0.5 μM Rubisco or 5 μM CsoS1A and CsoS1B and incubated for 15 min. Final buffer composition was 50 mM Tris, 150 mM NaCl, pH 7.5. NativeMark^TM^ Unstained Protein (life technologies) was used as the Mw marker.

### Cryo-EM of CsoSCA-Rubisco complex

0.5 μM Rubisco-strep was mixed with 0.5 mM of NTD_1-50_ peptide (CsoSCA residue 1-50) in 25 mM Tris pH 7.5, 80 mM NaCl containing 2% glycerol, incubated for 20 min at room temperature and thereafter stored on ice. 3.5 μL of this sample was deposited onto freshly glow-discharged (PELCO easiGlow), Quantifoil R 1.2/1.3 200 mesh Copper TEM grids (Quantifoil Microtools) and blotted for 3 sec using a Mark IV Vitrobot (FEI) after a 30 second delay under 100% humidity at 4 °C conditions before freezing in liquid ethane. The complex was visualized in a Talos Arctica (Thermo Fisher Scientific) operating at 200keV, and equipped with a K3 Summit director electron detector (Gatan) in super-resolution CDS mode at 57,000x, corresponding to a pixel size of 0.69 Å. In total, 5,742 movies were acquired with the aid of SerialEM * using a defocus range between -0.6 to -1.8 μm and a 3 x 3 multishot image shift pattern. All movies consisted of 50 frames with a total dose of 50 e-/Å^2^. The data collection was monitored using on-the-fly processing in cryoSPARC live (Structura Biotechnology Inc., https://cryosparc.com/live) (59) to monitor microscope performance, micrograph quality, and orientation distribution of the particles on the grid

### Image Processing

Super-resolution electron micrograph movies were aligned using MOTIONCOR2 (60) from within RELION 3.1 or using the CPU implementation of motion correction within RELION 3.1. CTF estimation was performed using CTFFIND 4.1 (61) from within RELION 3.1. Micrographs were inspected to remove poor-quality images, resulting in the higher quality selection of 3,932 micrographs. All further processing was done from within RELION 3.1.

Laplacian-of-Gaussian auto-picking was used on a subset of 200 micrographs to pick approximately 75,000 particles. These particles were extracted from the micrographs with a pixel size of 2.77 Ångstrom and a box size of 90 pixels. 2D classification was then used to generate a higher quality subset of particles that were used to generate an initial 3D model by way of Stochastic Gradient Descent. 3D classification of this higher quality subset of particles gave us a good quality 3D reference that was then used as a 3D template.

Approximately 1,300,000 particles were picked from the 3932 micrographs using our 3D reference before being extracted with a pixel size of 2.77 and a box size of 90 pixels. The particles were subjected to 3D classification applying D4 symmetry and a soft circular mask and the best-looking classes comprised of 358,785 particles were selected. The particles were then re-extracted with a pixel size of 1.37 Ångstrom and a box size of 180 pixels before undergoing another round of 3D classification. Again, the best classes comprised of 290,762 particles were selected. The particles were re-extracted with a pixel size of 0.91 Ångstrom and a box size of 312 pixels before being subjected to 3D auto-refinement. The refined particles were then 3D classified without additional image alignment and the best classes comprised of 262,882 particles were selected. These particles underwent CTF refinement and Bayesian polishing before being extracted with a larger 410 pixel size box. A few more rounds of 3D classification and 3D refinement, while selecting only the best classes left us with a homogeneous set of 79,562 particles. 3D refinement of this particle set gives a final resolution of 1.98 Ångstom at Fourier shell correlation (FSC) = 0.143. RELION 3.1 reports a b-factor of about -36 Å^2^ when sharpened with a soft mask.

### Coordinate model building and refinement

The coordinate models for the two *H. neapolitanus* Rubisco-NTD_1-50_ complex maps were built and refined similarly using a combination of COOT-v0.9.1 (62) and PHENIX-v1.19.1-4122 (63). Maps for this process were obtained by combination of the respective half-maps without filtering. Maps were molecular weight-based density modified and sharpened with *phenix.resolve_cryo_em* and *phenix.auto_sharpen* (64, 65), respectively. For ease of handling, the maps were reboxed to 160 vx^3^ (about 145 Å^3^) for further use. Chains A (CbbL) and D (CbbS) from the *Hnea* Rubisco-CsoS2 N*-peptide co-crystal structure (PDB ID: 6UEW) (29), stripped of all ligands, were used as initial models for both maps. The initial models were rigid-body docked and manually reworked to fit the maps in COOT and the resolved portion of CsoSCA NTD_1-50_ peptide built *de novo*. An initial round of *phenix.real_space_refinement* (66) was performed on models consisting of all asymmetric units with NCS constraints enforced, as well as default target bond length and angle restraints, but without secondary structure, rotamer, nor Ramachandran restraints. Putative ordered water molecules were then placed interactively in COOT using maps thresholded at 2σ based on presence of at least 2 hydrogen bonding partners and the occurrence of the density in both half-maps. Additional rounds of *phenix.real_space_refinement* and manual adjustment in COOT were performed as described above to yield the final coordinate models. For the higher resolution map (State-1), residues V3-E457 of CbbL were modelled with residues V324-E329 truncated to the Cᵦ atoms due to poor side-chain density in this region. Similarly, for the lower resolution map (State-2), residues V3-E457 of CbbL were modelled with residues H291-H300 and V323-D331 truncated to the Cᵦ atoms. The CbbS and NTD_1-50_ peptide density for both maps were modelled with residues M4-N110 and P22-A30, respectively.

### Visualization and structural analysis

Structural figures were prepared using a combination of PyMOL-v2.5 (Schrödinger, LLC.) and ChimeraX-v1.2.1 (67). Interface analysis to identify interacting residues and to calculate buried surface area was performed using the ePISA-v1.52 web server (68).

### Condensate formation assays

#### Labeling of CsoSCA-MBP

His-purified CsoSCA-MBP-his was run on a Superose 6 Increase 10/300 GL size exclusion column (GE Healthcare) to remove MBP-his contamination. Protein was eluted in 50 mM HEPES, 300 mM NaCl at pH 8. Aliquots containing CsoSCA-MBP were pooled. The pooled, dilute protein was labeled with Alexa Fluor 647 NHS Ester dye at a 1:1 ratio of protein to dye. Conjugation occurred over 2 hours in the dark at 4 °C. Excess dye was removed via buffer exchange into 50 mM HEPES, 300 mM NaCl on a EconoPac column (BioRad) and concentrated on a 30K cutoff spin column (Thermo Pierce) at 3500 x *g* for 20 minutes. Glycerol was added to a final concentration of 10% before flash-freezing the protein. *Condensate formation:* All condensate formation experiments were carried out in a final buffer concentration of 50 mM Tris, 20 mM NaCl, pH 7.5. The final concentrations were as follows: 1 μM Rubisco, 1 μM CsoS2-NTD-sfGFP, and 0.5 μM CsoSCA-MBP. For each experiment, the amount of carryover salt from the individual protein components was calculated to achieve a final concentration of 20 mM NaCl. 20 μl of each mixture was loaded onto a gasket fixed to a cover slip (CoverWell Perfusion Chamber 8×9 mm Dia × 0.9 mm Depth, Grace Bio-Labs) and imaged at 100x on a Zeiss Axio Observer Z1 inverted phase contrast microscope. Green channel excitation was 488 and emission was 509. Red channel excitation was 650 and emission was 673.

### Data availability

Cryo-EM maps (sharpened, full, and unfiltered halves) and masks have been deposited with the Electron Microscopy Data Bank, and the corresponding atomic coordinate models deposited with the Protein Data Bank for Rubisco-N50 peptide State-1 (EMD-25201, PDB-7SMK) and State-2 (EMD-25228, PDB-7SNV).

## Acknowledgment

We thank Paul Tobias for computational systems support and Luis Valentin for bioinformatics advice. We are grateful to Sacha Pulsford, Benedict Long, Avi Flamholz and Luke Oltrogge for advice and helpful discussion regarding the project and manuscript. Structural data were collected at Cal-Cryo at the California Institute for Quantitative Biosciences (QB3) of the University of California Berkeley. Molecular graphics and analyses was performed in part with UCSF ChimeraX, developed by the Resource for Biocomputing, Visualization, and Informatics at the University of California, San Francisco, with support from National Institutes of Health R01-GM129325 and the Office of Cyber Infrastructure and Computational Biology, National Institute of Allergy and Infectious Diseases. C.B. was partially supported by an International Postdoctoral grant from the Swedish Research Council (637-2014-6914). T.L. was supported by a NSF Graduate Research Fellowship (DGE 1106400). This work was funded by a grant from the U.S. Department of Energy (DE-SC00016240) to D.F.S.

## Author contributions

C.B. and D.F.S. designed the research. C.B., E.D. T.G.L. J.B, M.D.L, S.R.S and N.V. conducted the experiments. J.P.R. and T.G.L. solved the structure. C.B., T.G.L and D.F.S. wrote the manuscript with input and comments from all authors.

---

**SI Figure 1:**
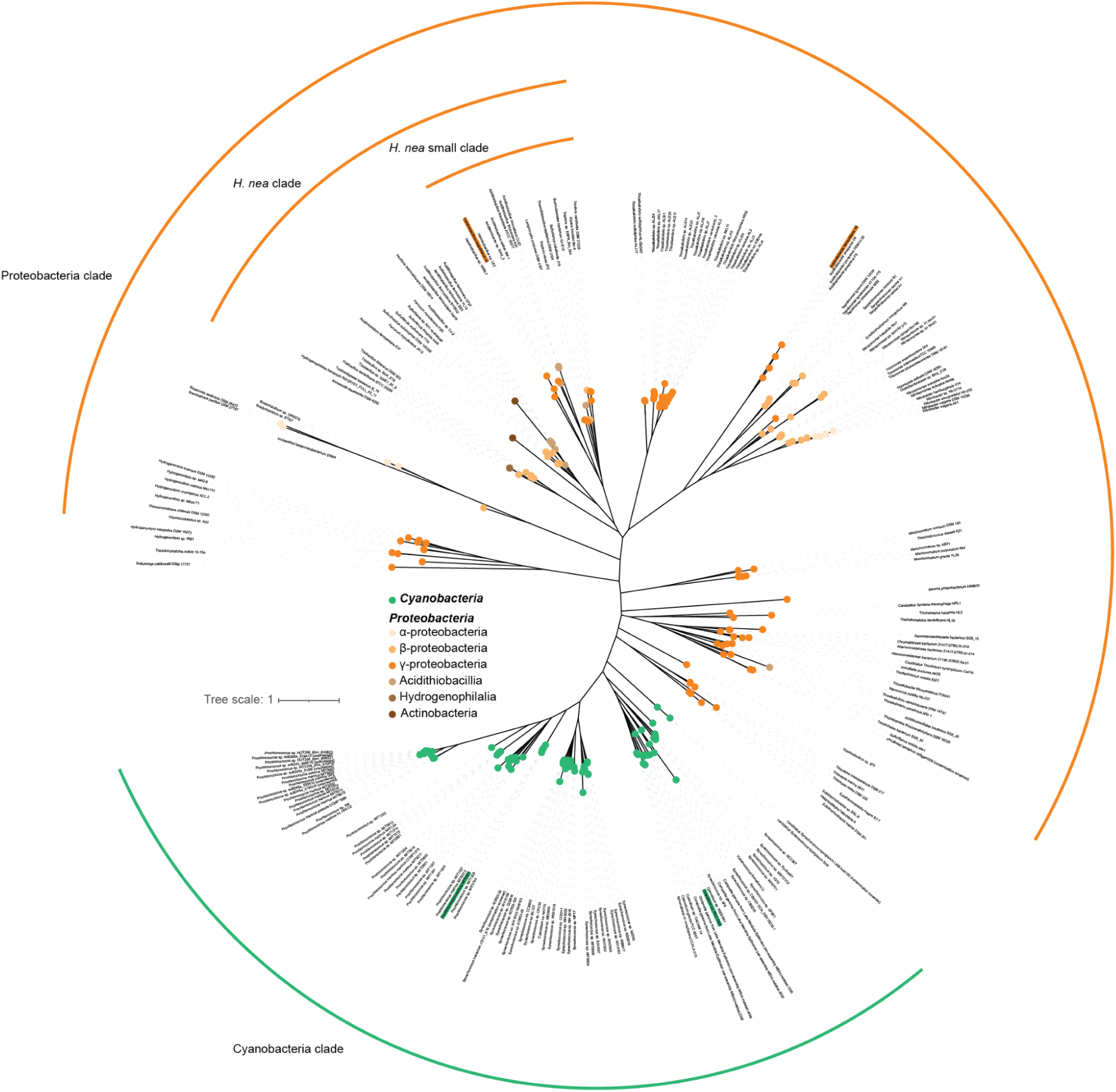
Phylogenetic tree of CsoSCA. Annotated maximum-likelihood phylogenetic tree of CsoSCA. Cyanobacterial homologs are colored in green and proteobacteria homologous in an orange/brown gradient. Scale bar, 0.1 substitutions per site.

**SI Figure 2:**
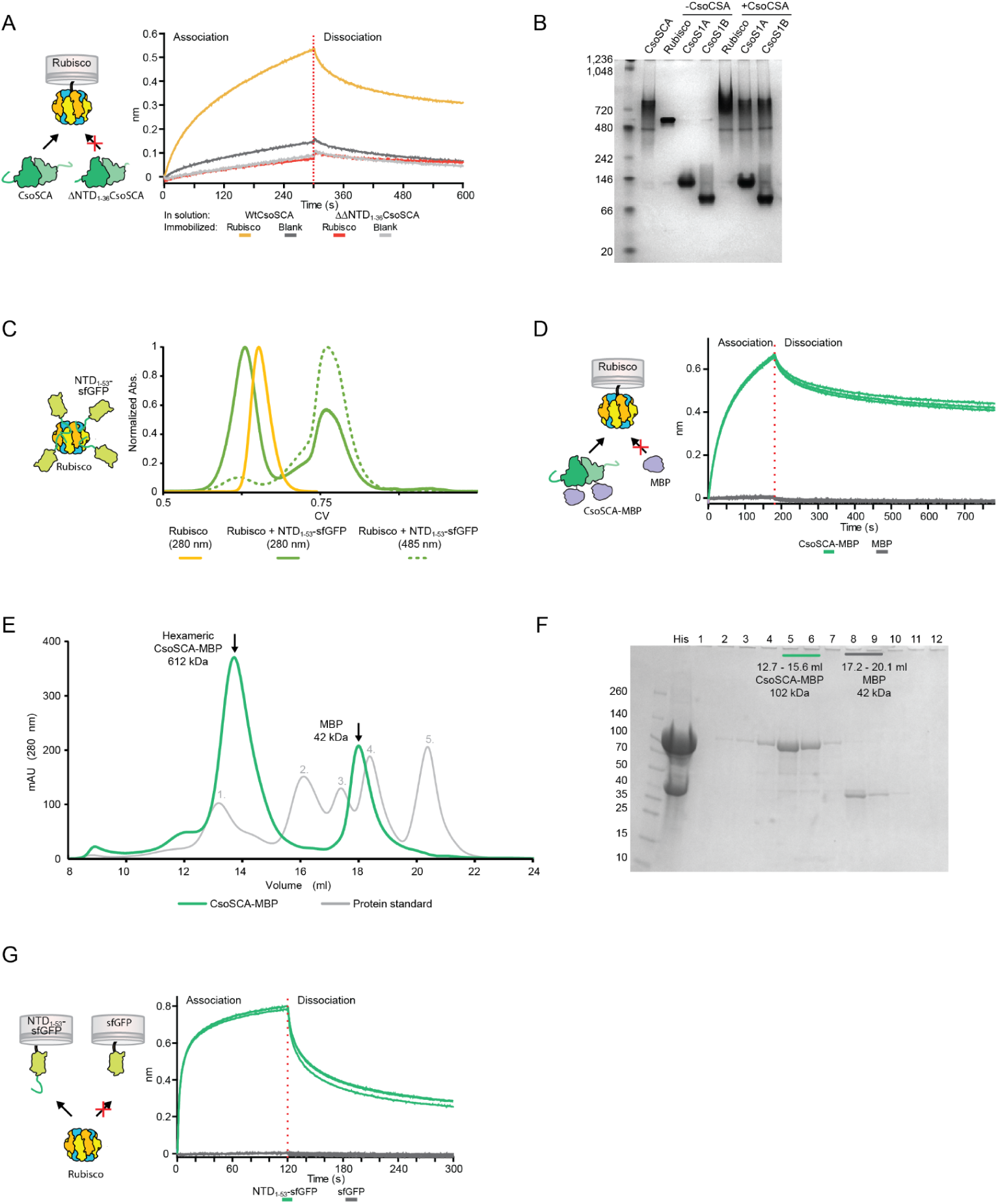
CsoSCA-Rubisco interaction controls. (A) BLI response showing that full length CsoSCA binds to Rubisco but a NTD truncated version (ΔNTD_1-37_CsoSCA) does not bind. (B) Native-page demonstrating binding between Rubisco and CsoSCA-MBP, and lack of binding between the major shell proteins CsoS1A and CsoS1B. Experiments were repeated in triplicate. (C) Size exclusion chromatography experiment showing co-elution of Rubisco and NTD_1-53_-sfGFP, demonstrating the interaction in a solution based assay. (D) BLI response showing that CsoSCA-MBP binds to Rubisco but not the negative control MBP. (E) Purification and determination of the oligomeric state of CsoSCA-MBP. Size exclusion-chromatogram of His-purified CsoSCA-MBP and Bio-Rad calibration standard. This is the second step of the purification and removes MBP-His contamination resulting in a pure CsoSCA-MBP complex. Expected molecular weight of the complex was determined to ∼600 kDa, indicating a trimer of dimer oligomeric state of CsoSCA-MBP. Molecular weight of standard was as follows: 1. Thyroglobin (670 kDa) 2. γ-globulin (158 kDa) 3. Ovalbumin (44 kDa) 3. Myoglobin (17 kDa) Vitamin B12 (1.4 kDa). (F) SDS-page gel on fractions from Size exclusion-chromatogram of His-purified CsoSCA-MBP. Marked fractions correspond to pure CsoSCA-MBP (Green, #5-6) and MBP (Grey, #8-9) peaks on chromatogram in (E). (G) BLI response showing that Rubisco binds to NTD_1-53_-sfGFP but not to the negative control sfGFP.

**SI Figure 3:**
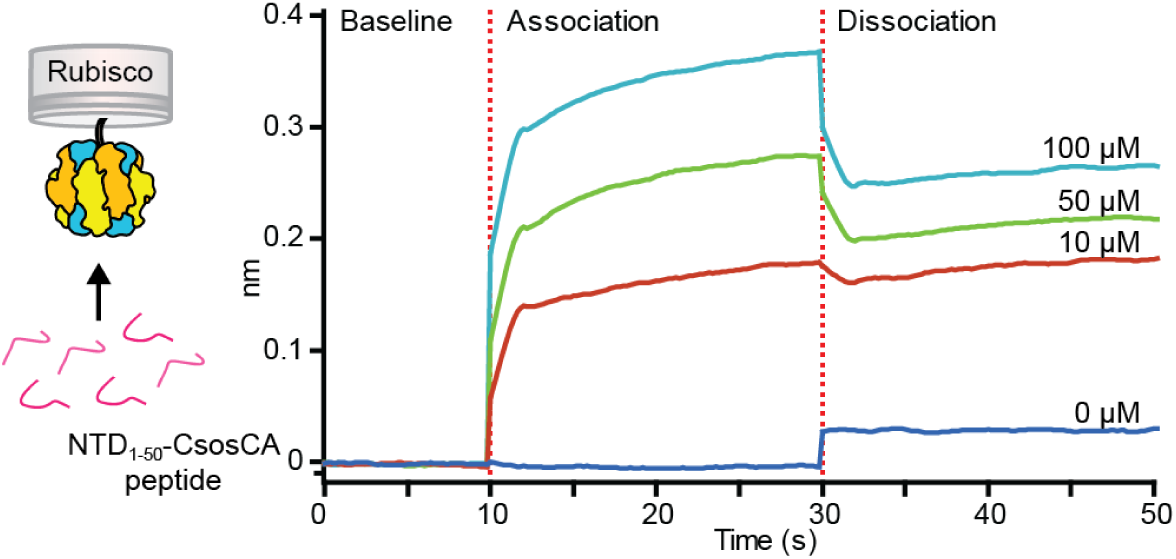
Binding of the NTD_1-50_CsoSCA peptide to immobilized Rubisco. BLI response showing binding of CsoSCA’s NTD_1-50_ peptide to Rubisco. Binding was measured with three peptide concentrations (10, 50 and 100 μM peptide).

**SI Figure 4:**
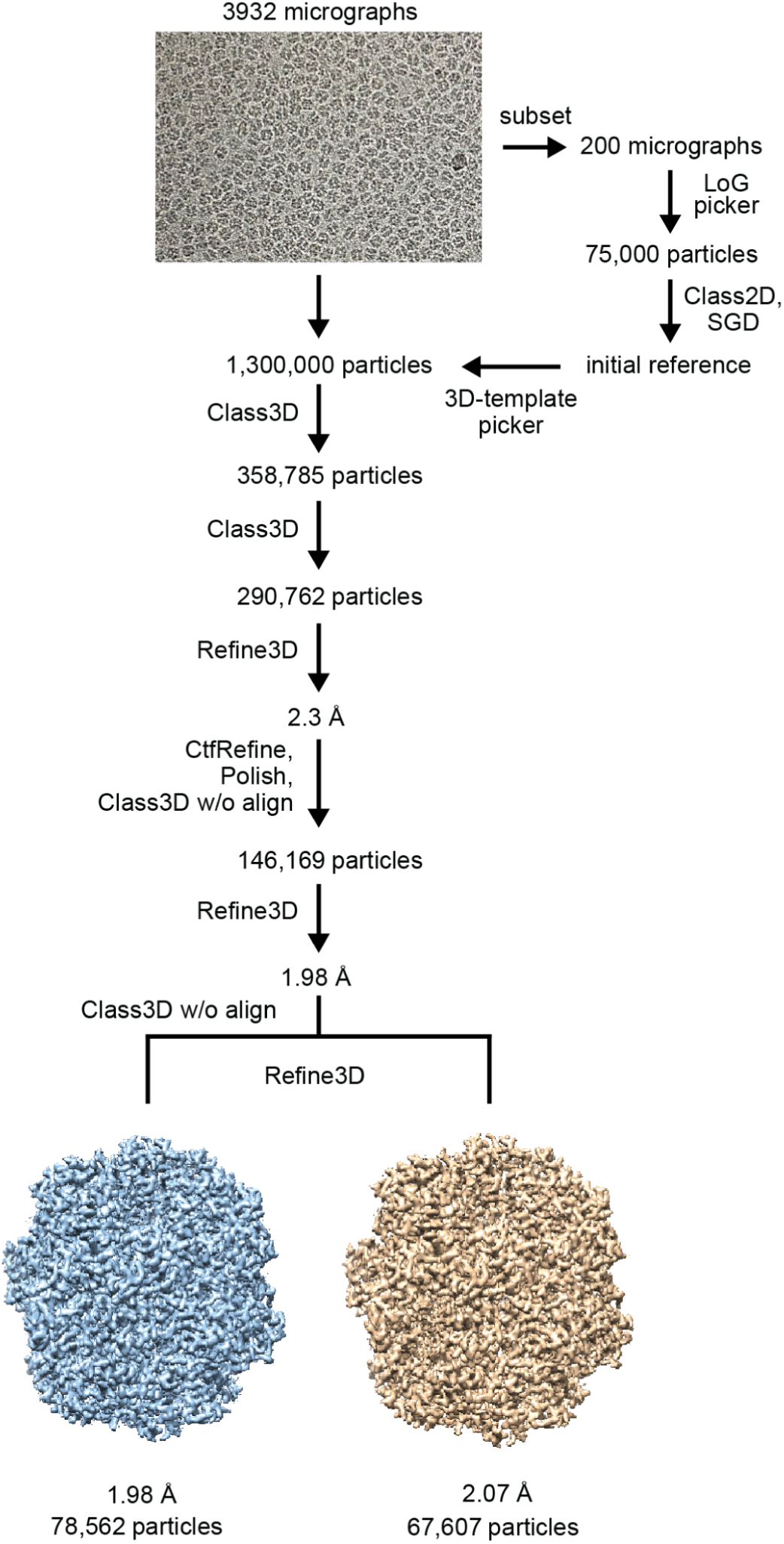
Single-particle Cryo-EM data collection and processing workflow. Representative micrograph of the Rubisco-NTD_1-50_CsoSCA complex and schematic of pre/processing, classification and refinement procedures used to generate the obtained in this study *see Methods for details).

**SI Figure 5:**
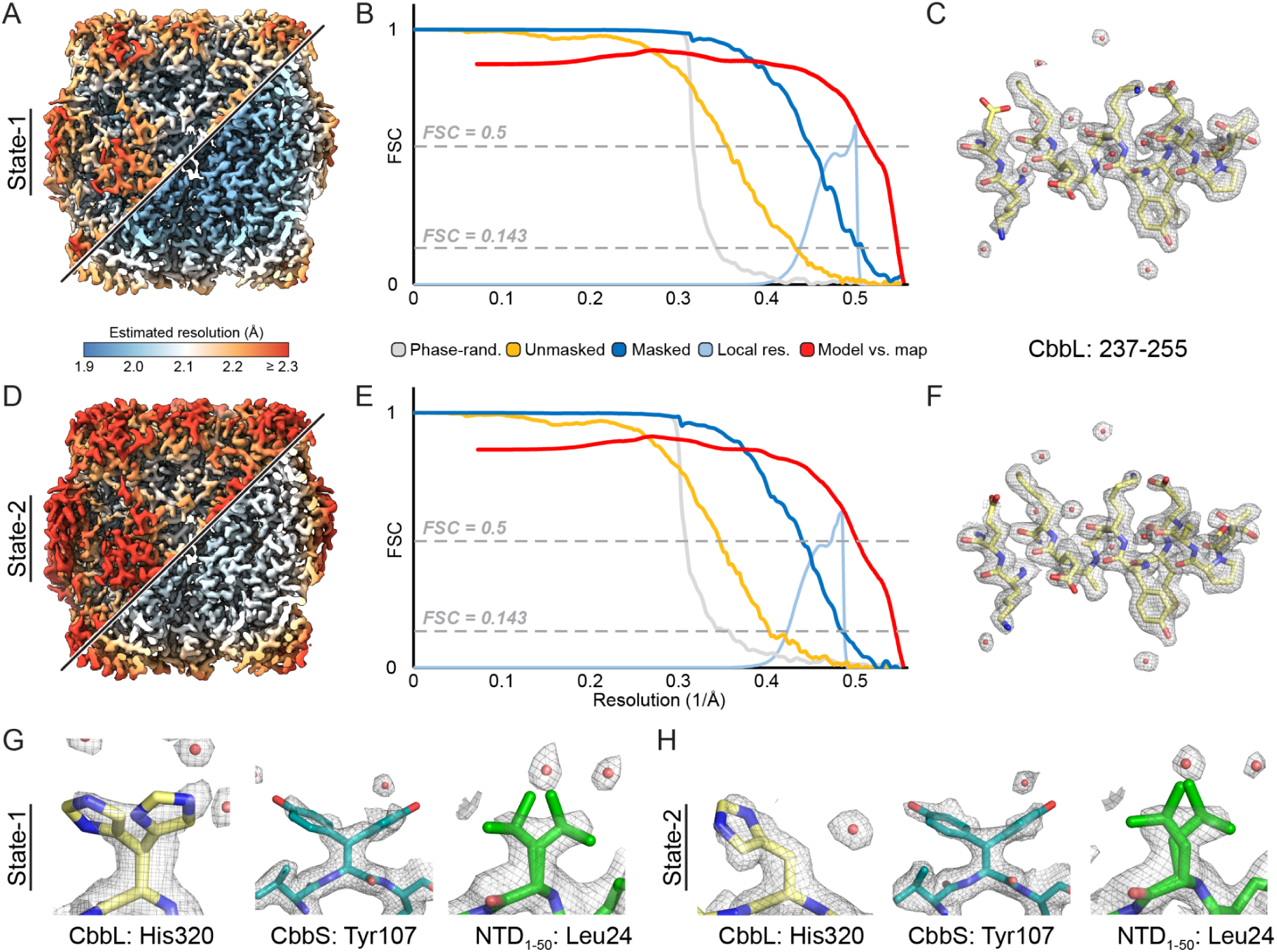
Resolution estimates and visual quality of Rubisco-NTD_1-50_CsoSCA complex cryo-EM reconstructions. (A) Local resolution estimate of State-1 map. The map is shown in unsharpened and locally filtered by estimated resolution. (B) Half-map and map-model FSC curves for State-1 map. (C) Exemplar State-1 density of an internal helix of CbbL from the density-modified and sharpened map contoured at 2σ. (D-F) Same as A-C for the State-2 cryo-EM reconstruction. (G,H) Model and density for residues with resolved alternate rotamer confirmations in either State-1 or State-2 reconstructions. Density shown is from the respective density-modified and sharpened map contoured at 2σ.

**SI Figure 6:**
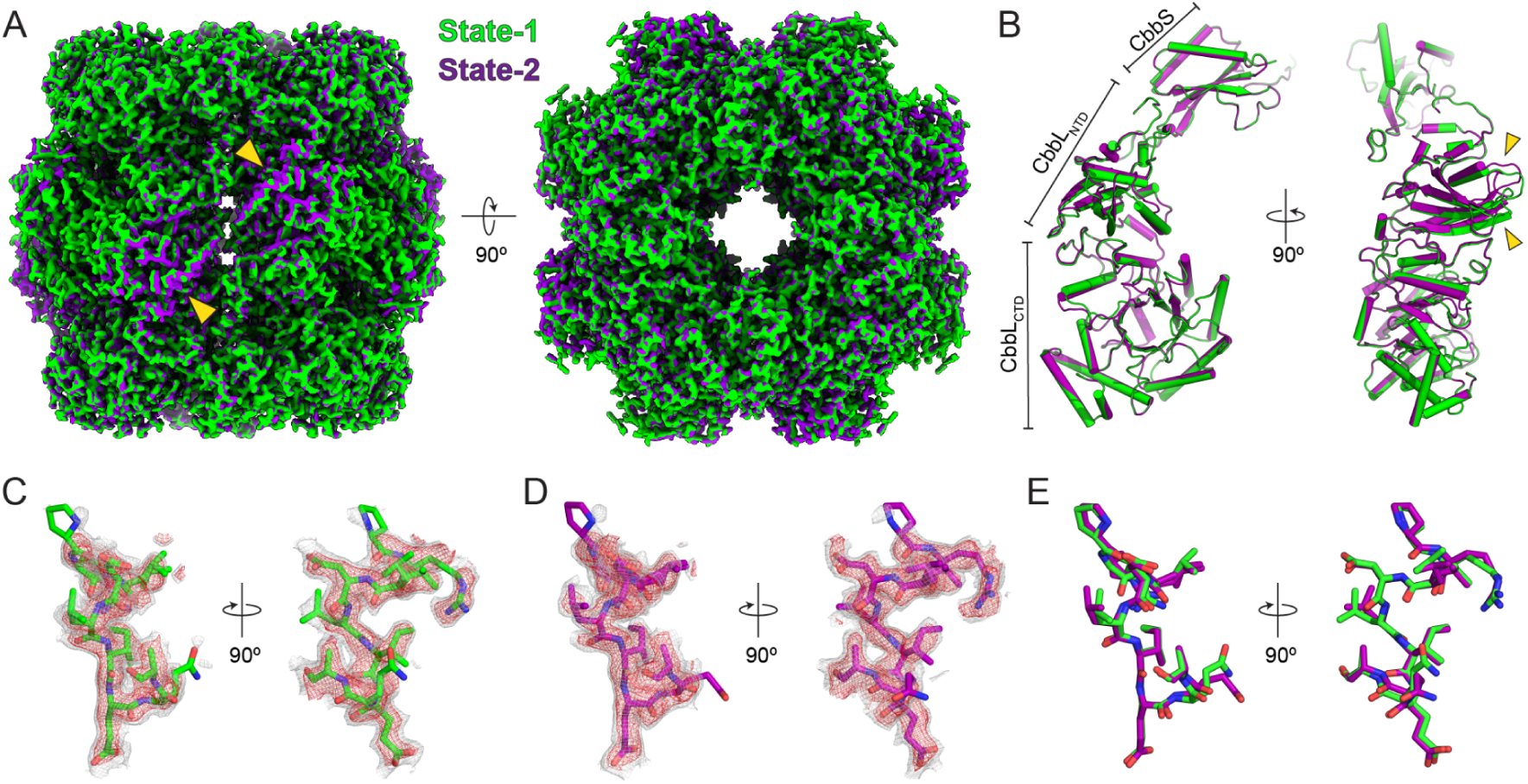
Comparison of the two reconstructed Rubisco-NTD_1-50_CsoSCA complex states. (A) Overlay of the two cryo-EM maps with State-1 in green and State-2 in purple. Yellow arrows point to regions of noticeable difference. (B) Asymmetric units of coordinate models for the two states, colored the same as in A. Subunits and domains are labeled and yellow arrows point to differences in loop confirmation in the CbbL_NTD_. (C) State-1 NTD_1-5_ peptide coordinate model and density contoured at 1.5σ (grey) and 2σ (red) from the density-modified, sharpened map. (D) Same as in C for State-2. (E) Overlay of NTD_1-50_ peptide coordinate models of the two states.

**SI Figure 7:**
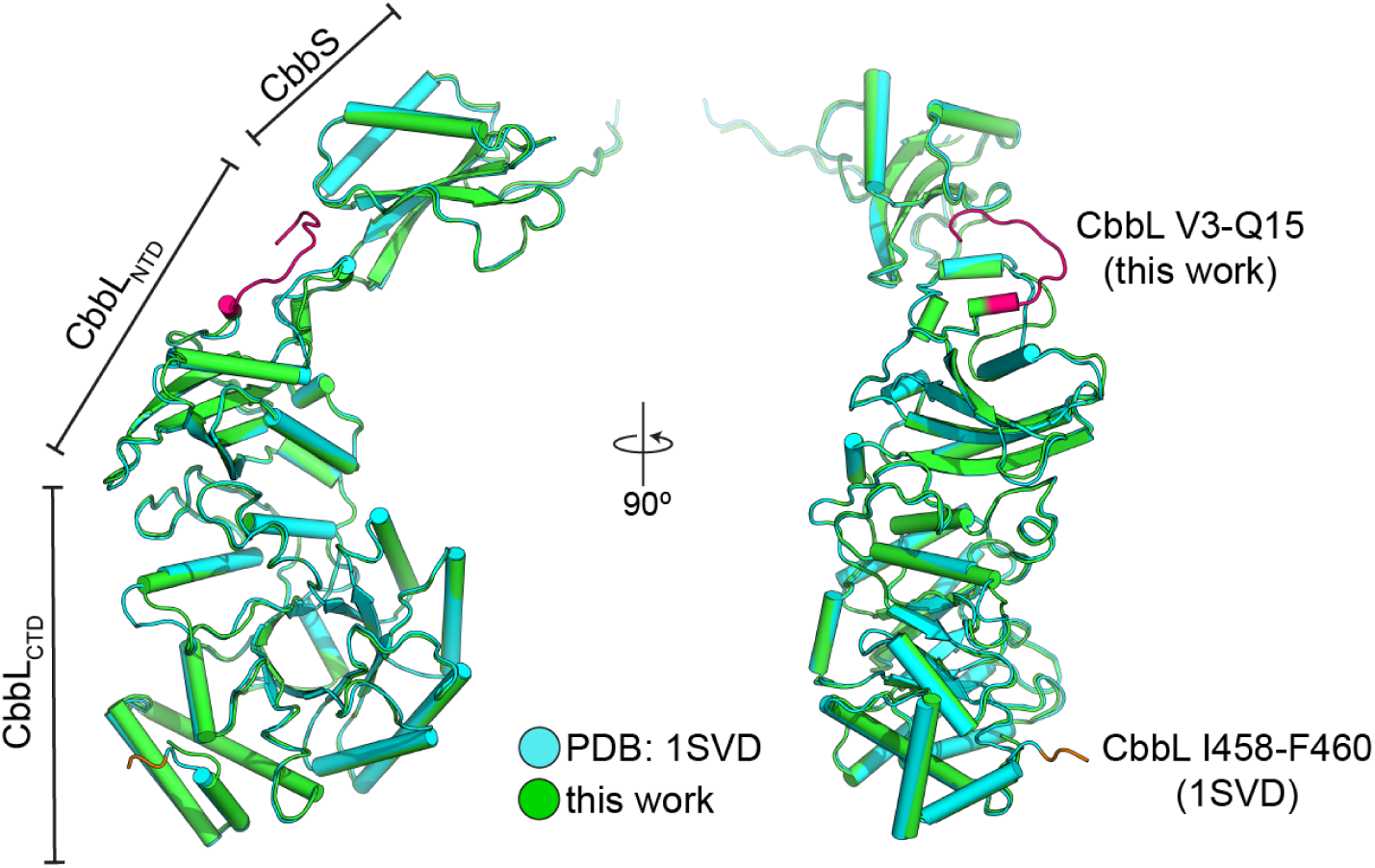
Comparison of cryo-EM and crystal structures of Hnea CbbL/S. Overlay of CbbL/S from State-1 (this work) in green and the previous X-ray crystal structure (PDB:1SVD) in cyan. Distinct subunits and domains are labeled. Differences in the extent of resolved termini of CbbL between these structures are highlighted. Magenta coloring indicates additional N-terminal CbbL residues resolved in this work. Orange coloring indicates C-terminal CbbL residues resolved in the crystal structure.

**SI Figure 8:**
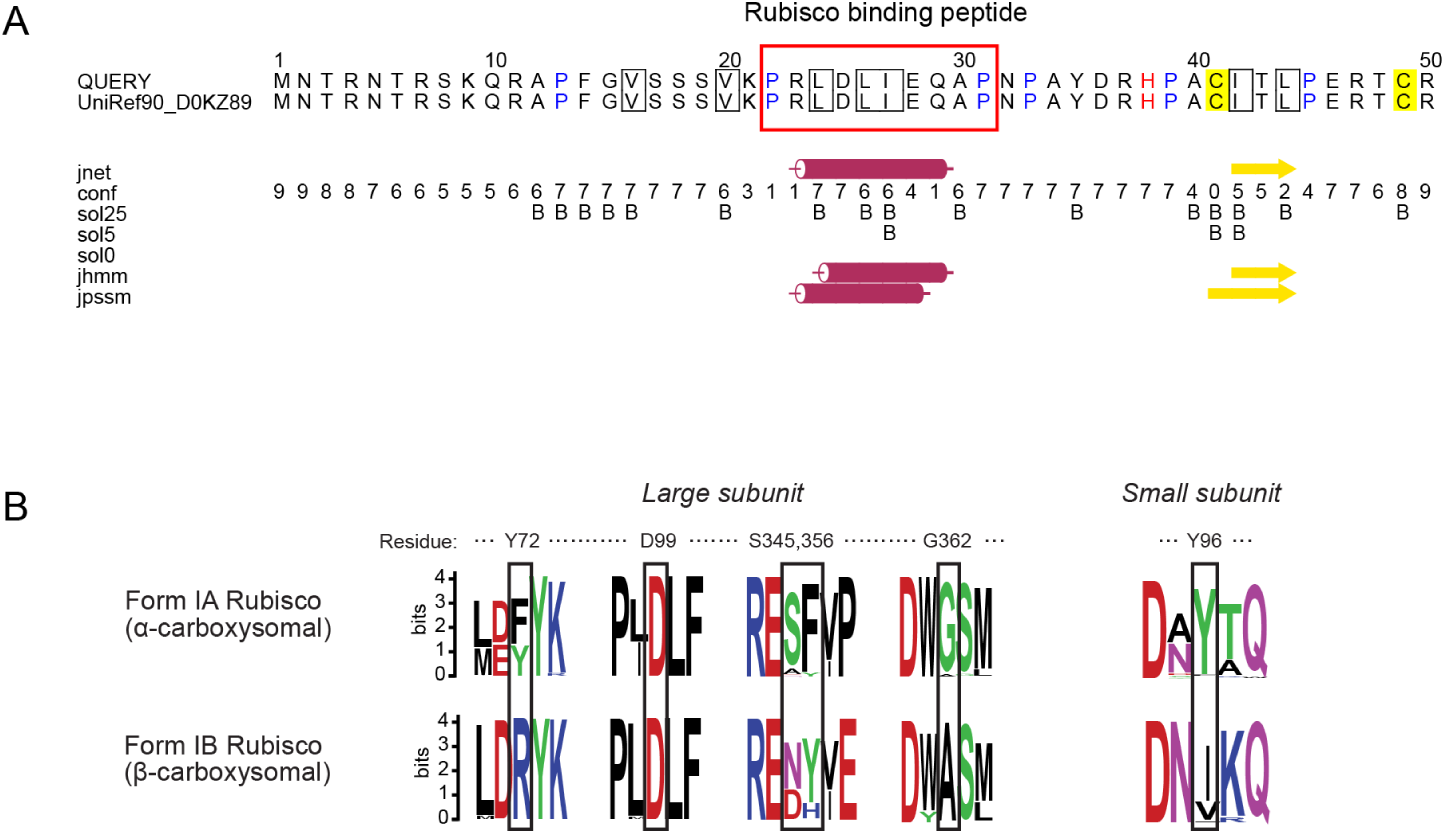
Secondary structure prediction of NTD_1-50_CsoSCA peptide and Rubisco sequence comparison. (A) JPred4 protein secondary structure prediction server was used to predict the secondary structure of *H. neapolitanus* CsoSCA’s NTD_1-50_ sequence. The PRLDLIEQAP sequence is predicted to form an alpha-helical structure (red cylinder). This stretch of sequence corresponds to the extra density observed in the cryo-EM structure of Rubisco in complex with the NTD_1-50_ peptide (red box). (B) Rubisco sequence comparison at the CsoSCA-peptide interaction site. Multiple sequence alignment of α-carboxysomal Form IA Rubiscos and β-carboxysomal Form IB Rubiscos visualized using Weblogo (SI Table 1). Residues are numbered according to the Rubisco *H. neapolitanus* sequence. Residues shown in our Rubisco *H. neapolitanus* structure to interact with the CsoSCA peptide are marked with a black box. CsoSCA interacting residues have a high conservation score but are, in general, not conserved between α-carboxysomal and β-carboxysomal Rubiscos.

**SI Figure 9:**
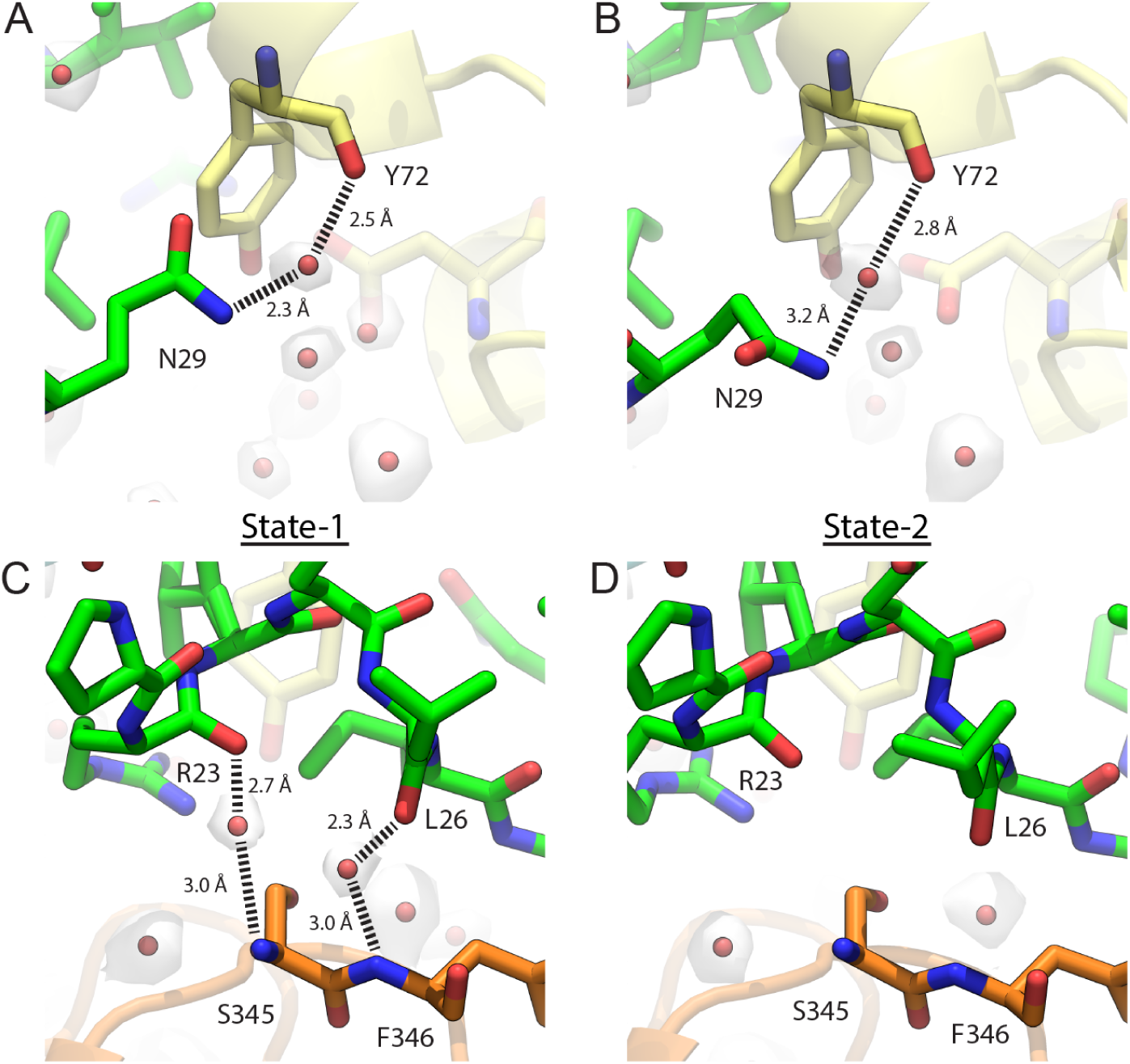
Putative ordered waters mediating interaction between NTD_1-50_CsoSCA peptide and Rubisco. (A,B) Water-mediated interaction between CsoSCA’s NTD_1-50_ peptide and CbbL_A_ resolved in both Rubisco-NTD_1-50_. (C,D) Water-mediated interactions between NTD_1-50_ peptide and CbbL_B_ resolved in the State-1 (C, 1.98 Å) map but not in State-2 (D, 2.07 Å) map. Putative ordered water densities are shown as transparent white surfaces contoured at 1.5σ from respective density-modified and sharpened maps.

**SI Figure 10:**
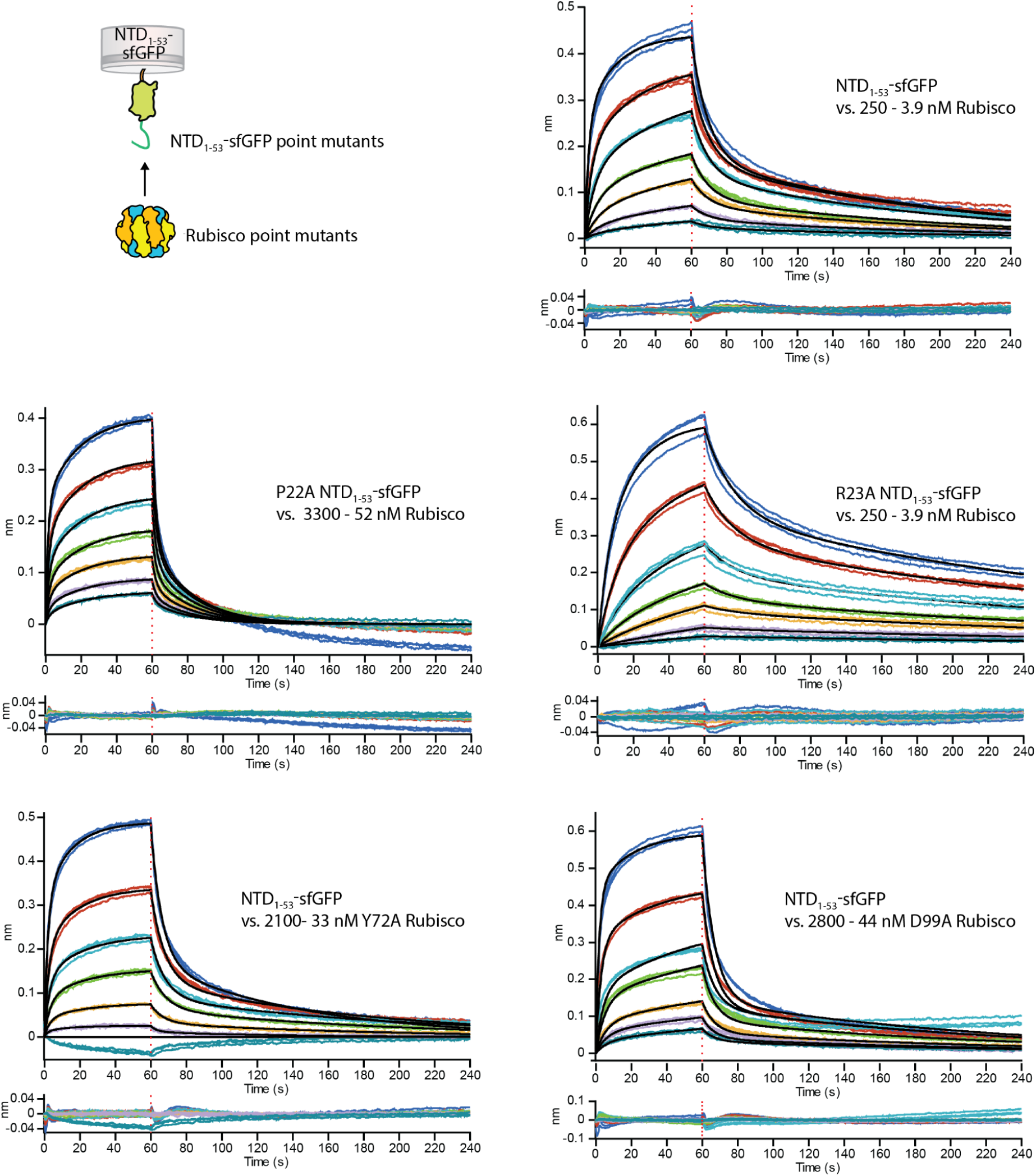
BLI sensograms of NTD_1-53_-sfGFP and Rubisco point mutants. BLI response from binding affinity measurements of Rubisco against immobilized NTD_1-53_-sfGFP using point mutants of Rubisco or of NTD_1-53_-sfGFP. Mutant and concentration range of Rubisco are indicated in the figure. *K*_D_, *k*_on_ and *k*_off_ are listed in Table 1 and SI Table 4.

**SI Figure 11:**
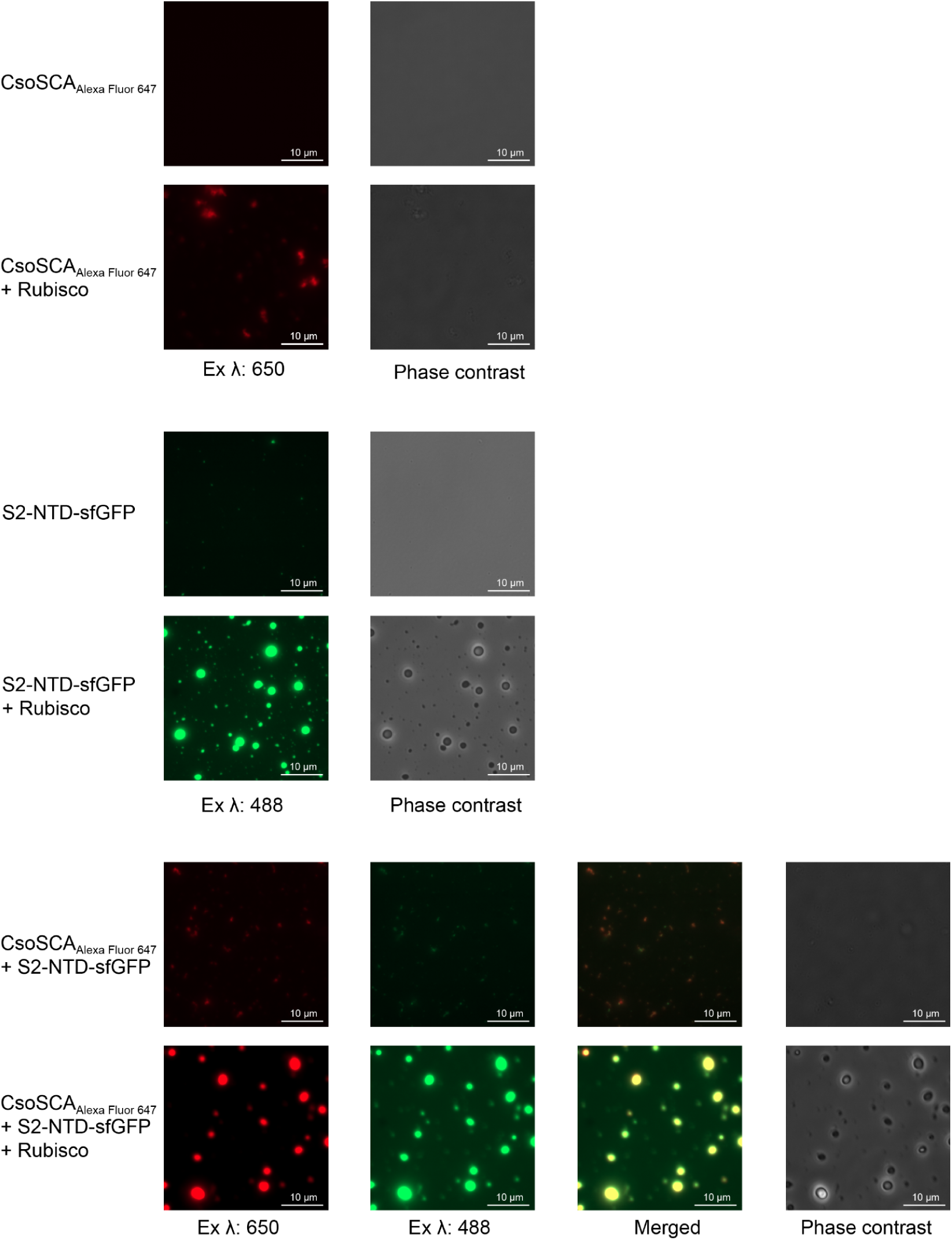
Condensate formation with CsoS2, Rubisco and CsoSCA. Alexa Fluor 647, sfGFP and merged fluorescence as well as phase contrast images of protein droplets formed from a solution of Rubisco, CsoS2-NTD-sfGFP and Alexa Fluor 647 labeled CsoSCA-MBP (CsoSCA_Alexa Fluor 647_) and of negative controls. Micrographs show that CsoSCA recruits into Rubisco-CsoS2-NTD protein condensates.

**SI Figure 12:**
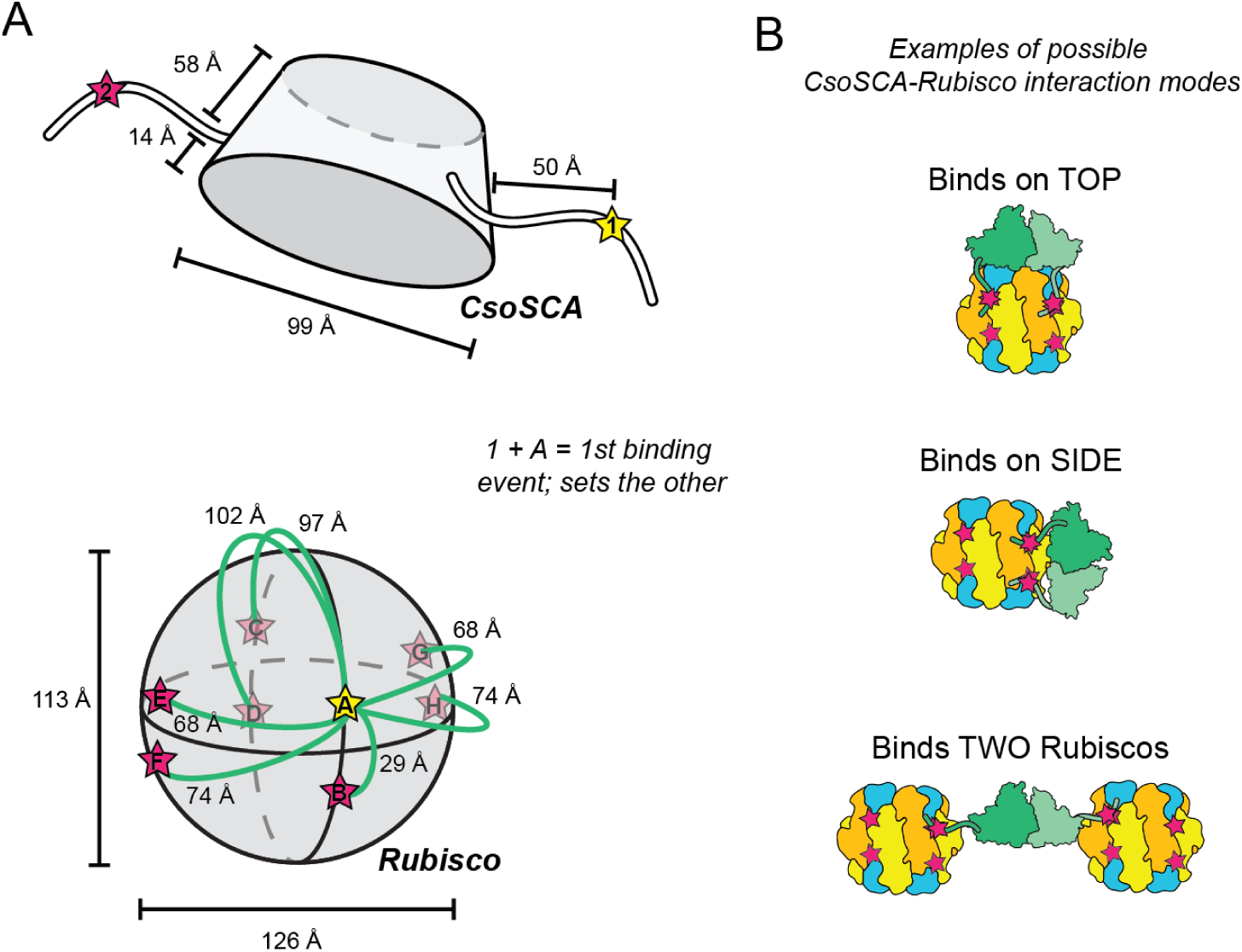
Possible binding conformations of CsoSCA-Rubisco. (A) Diagram showing the approximate distances between binding motif on CsoSCA (top) and binding sites on Rubisco (bottom). Binding motifs/sites are marked with a star. (B) Cartoon representation of three examples of possible CsoSCA-Rubisco interaction modes.

**SI Figure 13:**
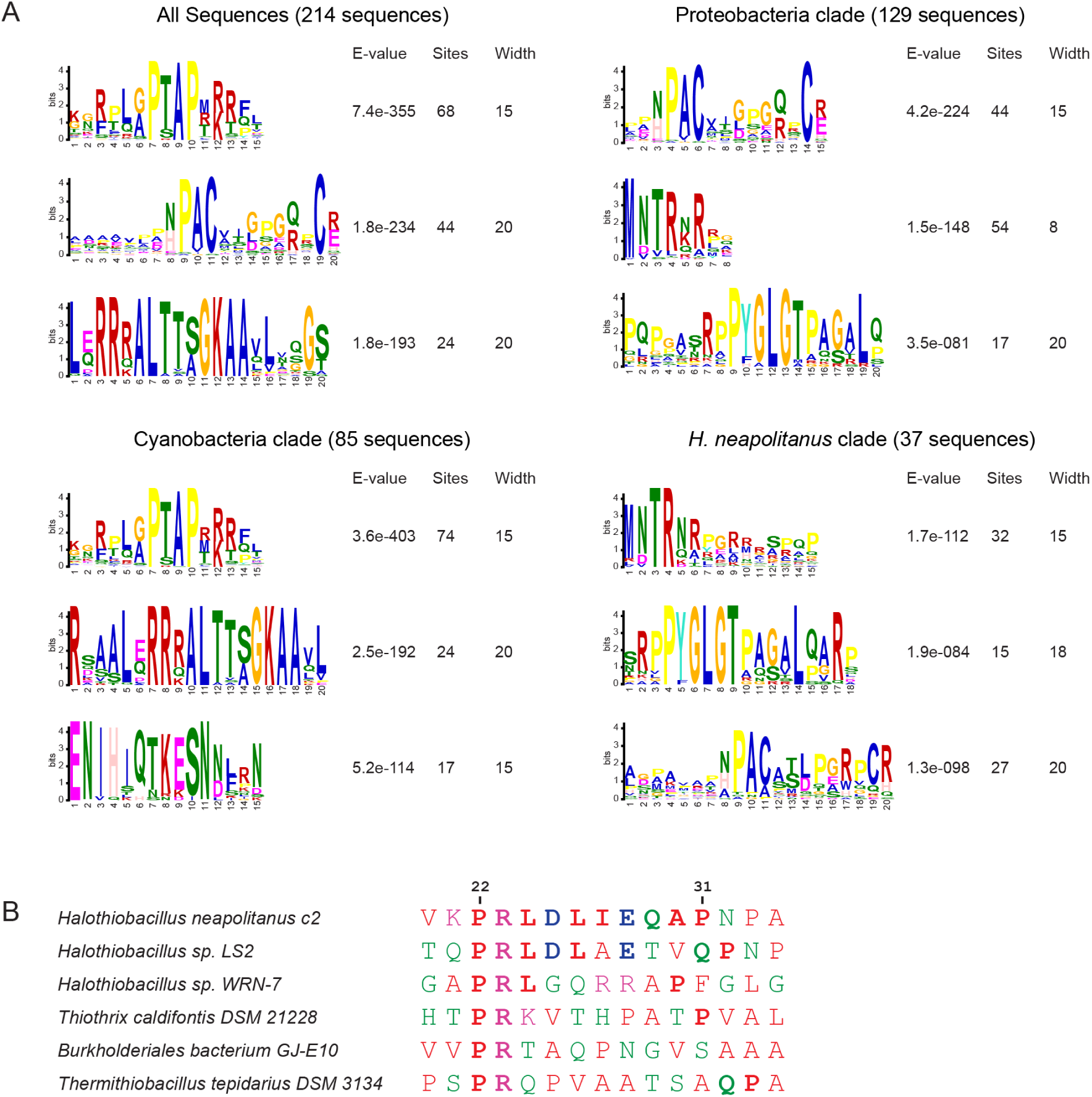
Conservation analysis of CsoSCA NTD Rubisco binding motif. (A) Motif discovery of CsoSCAs NTD sequence using MEME analysis (Multiple Em for Motif Elicitation). Analysis was performed on all CsoSCA sequences, as well as sequences specific to the proteobacteria and cyanobacteria (SI Table 1). For a more stringent analysis, the *H. neapolitanus* clade was analysed separately as well. 44/129 of the collected proteobacteria sequences contain a PACxxxxxxxC motif. *H. neapolitanus* CsoSCA has this motif, however, it does not appear to be essential for binding to Rubisco. The N-terminus MNTRxR is somewhat conserved in proteobacteria (55/129 sequences). In cyanobacterial CsoSCAs a conserved PTAPxRR motif (75/85 sequences) is identified. 24/85 cyanobacteria sequences contain what appears to be the conserved CsoS2 Rubisco binding motif, RxxxxxRRRxxxxxGK. These sequences all belong to the *Procloroccocos* genus. The experimentally identified *H. neapolitanus* CsoSCA Rubisco binding motif, PRLDLIEQA, does not appear to be conserved among homologos. (B) Alignment of a selection of sequences from *H. neapolitanus* CsoSCAs closest homologs. Bold letters indicate residues conserved with the *H. neapolitanus* motif. Its closest homologue *Halothiobacillus sp. LS2* contains the same motif. Some of the others have the PR residues, demonstrated to be essential for binding, however the rest of the motif is poorly conserved.

**SI Table 3:** Cryo-EM data collection, refinement and model statistics.

|  | *State 1 (ASU)*<br>(EMDB-25201)<br>(PDB 7SMK) | *State 2 (ASU)*<br>(EMDB-25228)<br>(PDB 7SNV) |
| --- | --- | --- |
| **Data collection and processing** |  |  |
| Microscope | TFS Talos Arctica |  |
| Detector | Gatan K3 |  |
| Voltage (kV) | 200 |  |
| Nominal magnification | 57,000 |  |
| Pixel size (Å) | 0.69 |  |
| Electron exposure (e-/Å<sup>2</sup>) | 50 |  |
| Defocus range (µm) | -0.6 to -1.8 |  |
| Symmetry imposed | D4 |  |
| Initial particle images (no.) | ~1.3 mil |  |
| Final particle images (no.) | 78,562 | 67,607 |
| Map resolution (Å) | 1.98 | 2.07 |
| FSC threshold | 0.143 |  |
| Map resolution range (Å) | 1.9-2.7 | 2-2.9 |
| **Coordinate model refinement** |  |  |
| Initial model used (PDB code) | 6UEW |  |
| Model resolution (Å) | 1.94 | 2.02 |
| FSC threshold | 0.5 |  |
| Map sharpening *B* factor (Å<sup>2</sup>) | *phenix.autosharpen* |  |
| Model composition |  |  |
| Non-hydrogen atoms | 4729 | 4685 |
| Protein residues | 571 | 571 |
| Ligands (waters) | 202 | 200 |
| Atomic displacement parameters (Å<sup>2</sup>) |  |  |
| Protein | 21.50 | 14.47 |
| Ligand (waters) | 22.68 | 13.10 |
| R.m.s. deviations |  |  |
| Bond lengths (Å) | 0.008 | 0.007 |
| Bond angles (°) | 1.156 | 1.061 |
| Validation |  |  |
| MolProbity score | 1.79 | 1.71 |
| Clashscore | 6.96 | 7.61 |
| Poor rotamers (%) | 2.15 | 0.88 |
| Ramachandran plot (%) |  |  |
| Favored | 97.17 | 95.75 |
| Allowed | 2.65 | 4.07 |
| Disallowed | 0.18 | 0.18 |

## References

1. C. T. Supuran, Carbonic anhydrases: novel therapeutic applications for inhibitors and activators. Nat. Rev. Drug Discov. 7, 168–181 (2008).

2. C. Merlin, M. Masters, S. McAteer, A. Coulson, Why is carbonic anhydrase essential to Escherichia coli? J. Bacteriol. 185, 6415–6424 (2003).

3. M. R. Badger, G. D. Price, The role of carbonic anhydrase in photosynthesis. Annu. Rev. Plant Physiol. Plant Mol. Biol. 45, 369–392 (1994).

4. I. Andersson, Catalysis and regulation in Rubisco. J. Exp. Bot. 59, 1555–1568 (2008).

5. C. Bathellier, et al., Ribulose 1,5-bisphosphate carboxylase/oxygenase activates O2 by electron transfer. Proc Natl Acad Sci USA 117, 24234–24242 (2020).

6. A. I. Flamholz, et al., Revisiting Trade-offs between Rubisco Kinetic Parameters. Biochemistry 58, 3365–3376 (2019).

7. A. Flamholz, P. M. Shih, Cell biology of photosynthesis over geologic time. Curr. Biol. 30, R490–R494 (2020).

8. C. A. Kerfeld, M. R. Melnicki, Assembly, function and evolution of cyanobacterial carboxysomes. Curr. Opin. Plant Biol. 31, 66–75 (2016).

9. B. D. Rae, B. M. Long, M. R. Badger, G. D. Price, Functions, compositions, and evolution of the two types of carboxysomes: polyhedral microcompartments that facilitate CO2 fixation in cyanobacteria and some proteobacteria. Microbiol. Mol. Biol. Rev. 77, 357–379 (2013).

10. J. J. Desmarais, et al., DABs are inorganic carbon pumps found throughout prokaryotic phyla. Nat. Microbiol. 4, 2204–2215 (2019).

11. A. I. Flamholz, et al., Functional reconstitution of a bacterial CO2 concentrating mechanism in Escherichia coli. eLife 9 (2020).

12. F. Cai, et al., The pentameric vertex proteins are necessary for the icosahedral carboxysome shell to function as a CO2 leakage barrier. PLoS ONE 4, e7521 (2009).

13. N. Mangan, M. Brenner, Systems analysis of the CO2 concentrating mechanism in cyanobacteria. eLife, e02043 (2014).

14. N. M. Mangan, A. Flamholz, R. D. Hood, R. Milo, D. F. Savage, pH determines the energetic efficiency of the cyanobacterial CO2 concentrating mechanism. Proc Natl Acad Sci USA 113, E5354–62 (2016).

15. M. R. Melnicki, M. Sutter, C. A. Kerfeld, Evolutionary relationships among shell proteins of carboxysomes and metabolosomes. Curr. Opin. Microbiol. 63, 1–9 (2021).

16. M. S. Kimber, Carboxysomal carbonic anhydrases. Subcell Biochem 75, 89–103 (2014).

17. Z. Dou, et al., CO2 fixation kinetics of Halothiobacillus neapolitanus mutant carboxysomes lacking carbonic anhydrase suggest the shell acts as a diffusional barrier for CO2. J. Biol. Chem. 283, 10377–10384 (2008).

18. H. Fukuzawa, E. Suzuki, Y. Komukai, S. Miyachi, A gene homologous to chloroplast carbonic anhydrase (icfA) is essential to photosynthetic carbon dioxide fixation by Synechococcus PCC7942. Proc Natl Acad Sci USA 89, 4437–4441 (1992).

19. G. D. Price, M. R. Badger, Expression of Human Carbonic Anhydrase in the Cyanobacterium Synechococcus PCC7942 Creates a High CO(2)-Requiring Phenotype: Evidence for a Central Role for Carboxysomes in the CO(2) Concentrating Mechanism. Plant Physiol. 91, 505–513 (1989).

20. S. H. Baker, D. S. Williams, H. C. Aldrich, A. C. Gambrell, J. M. Shively, Identification and localization of the carboxysome peptide Csos3 and its corresponding gene in Thiobacillus neapolitanus. Arch. Microbiol. 173, 278–283 (2000).

21. A. K.-C. So, et al., A novel evolutionary lineage of carbonic anhydrase (epsilon class) is a component of the carboxysome shell. J. Bacteriol. 186, 623–630 (2004).

22. M. R. Sawaya, et al., The structure of beta-carbonic anhydrase from the carboxysomal shell reveals a distinct subclass with one active site for the price of two. J. Biol. Chem. 281, 7546–7555 (2006).

23. S. Heinhorst, et al., Characterization of the carboxysomal carbonic anhydrase CsoSCA from Halothiobacillus neapolitanus. J. Bacteriol. 188, 8087–8094 (2006).

24. K. L. Peña, S. E. Castel, C. de Araujo, G. S. Espie, M. S. Kimber, Structural basis of the oxidative activation of the carboxysomal gamma-carbonic anhydrase, CcmM. Proc Natl Acad Sci USA 107, 2455–2460 (2010).

25. C. de Araujo, et al., Identification and characterization of a carboxysomal γ-carbonic anhydrase from the cyanobacterium Nostoc sp. PCC 7120. Photosyn. Res. 121, 135–150 (2014).

26. L. D. McGurn, et al., The structure, kinetics and interactions of the β-carboxysomal β-carbonic anhydrase, CcaA. Biochem. J. 473, 4559–4572 (2016).

27. B. M. Long, M. R. Badger, S. M. Whitney, G. D. Price, Analysis of carboxysomes from Synechococcus PCC7942 reveals multiple Rubisco complexes with carboxysomal proteins CcmM and CcaA. J. Biol. Chem. 282, 29323–29335 (2007).

28. K. Zang, H. Wang, F. U. Hartl, M. Hayer-Hartl, Scaffolding protein CcmM directs multiprotein phase separation in β-carboxysome biogenesis. Nat. Struct. Mol. Biol. 28, 909–922 (2021).

29. L. M. Oltrogge, et al., Multivalent interactions between CsoS2 and Rubisco mediate α-carboxysome formation. Nat. Struct. Mol. Biol. 27, 281–287 (2020).

30. S. He, et al., The structural basis of Rubisco phase separation in the pyrenoid. Nat. Plants 6, 1480–1490 (2020).

31. M. T. Meyer, et al., Assembly of the algal CO2-fixing organelle, the pyrenoid, is guided by a Rubisco-binding motif. Sci. Adv. 6 (2020).

32. M. Wu, G. C. Lander, M. A. Herzik, Sub-2 Angstrom resolution structure determination using single-particle cryo-EM at 200 keV. Journal of Structural Biology: X 4, 100020 (2020).

33. F. Cai, et al., Advances in understanding carboxysome assembly in prochlorococcus and synechococcus implicate csos2 as a critical component. Life (Basel) 5, 1141–1171 (2015).

34. T. Chaijarasphong, et al., Programmed Ribosomal Frameshifting Mediates Expression of the α-Carboxysome. J. Mol. Biol. 428, 153–164 (2016).

35. Y. Sun, et al., Decoding the absolute stoichiometric composition and structural plasticity of α-carboxysomes. BioRxiv (2021) https://doi.org/10.1101/2021.12.06.471529.

36. F. Cai, S. Heinhorst, J. M. Shively, G. C. Cannon, Transcript analysis of the Halothiobacillus neapolitanus cso operon. Arch. Microbiol. 189, 141–150 (2008).

37. V. N. Uversky, Intrinsically disordered proteins in overcrowded milieu: Membrane-less organelles, phase separation, and intrinsic disorder. Curr. Opin. Struct. Biol. 44, 18–30 (2017).

38. L. A. Metskas, et al., Rubisco forms a lattice inside alpha-carboxysomes. Nat. Commun. 13, 4863 (2022).

39. T. Ni, et al., Structure and assembly of cargo Rubisco in two native α-carboxysomes. Nat. Commun. 13, 4299 (2022).

40. Y.-C. C. Tsai, M. C. Lapina, S. Bhushan, O. Mueller-Cajar, Identification and characterization of multiple rubisco activases in chemoautotrophic bacteria. Nat. Commun. 6, 8883 (2015).

41. M. Sutter, et al., Structural Characterization of a Newly Identified Component of α-Carboxysomes: The AAA+ Domain Protein CsoCbbQ. Sci. Rep. 5, 16243 (2015).

42. H. Wang, et al., Rubisco condensate formation by CcmM in β-carboxysome biogenesis. Nature 566, 131–135 (2019).

43. P. Ryan, et al., The small RbcS-like domains of the β-carboxysome structural protein CcmM bind RubisCO at a site distinct from that binding the RbcS subunit. J. Biol. Chem. 294, 2593–2603 (2019).

44. S. Lechno-Yossef, et al., Cyanobacterial carboxysomes contain an unique rubisco-activase-like protein. New Phytol. 225, 793–806 (2020).

45. M. Flecken, et al., Dual functions of a rubisco activase in metabolic repair and recruitment to carboxysomes. Cell 183, 457–473.e20 (2020).

46. B. M. Long, B. Förster, S. B. Pulsford, G. D. Price, M. R. Badger, Rubisco proton production can drive the elevation of CO2 within condensates and carboxysomes. Proc Natl Acad Sci USA 118 (2021).

47. A. I. Flamholz, et al., Trajectories for the evolution of bacterial CO2-concentrating mechanisms. Proc Natl Acad Sci USA 119, e2210539119 (2022).

48. L. C. M. Mackinder, et al., A repeat protein links Rubisco to form the eukaryotic carbon-concentrating organelle. Proc Natl Acad Sci USA 113, 5958–5963 (2016).

49. T. Wunder, S. L. H. Cheng, S.-K. Lai, H.-Y. Li, O. Mueller-Cajar, The phase separation underlying the pyrenoid-based microalgal Rubisco supercharger. Nat. Commun. 9, 5076 (2018).

50. V. M. Markowitz, et al., IMG: the Integrated Microbial Genomes database and comparative analysis system. Nucleic Acids Res. 40, D115–22 (2012).

51. F. Madeira, et al., The EMBL-EBI search and sequence analysis tools APIs in 2019. Nucleic Acids Res. 47, W636–W641 (2019).

52. A. M. Waterhouse, J. B. Procter, D. M. A. Martin, M. Clamp, G. J. Barton, Jalview Version 2--a multiple sequence alignment editor and analysis workbench. Bioinformatics 25, 1189–1191 (2009).

53. J. Trifinopoulos, L.-T. Nguyen, A. von Haeseler, B. Q. Minh, W-IQ-TREE: a fast online phylogenetic tool for maximum likelihood analysis. Nucleic Acids Res. 44, W232–5 (2016).

54. I. Letunic, P. Bork, Interactive Tree Of Life (iTOL) v5: an online tool for phylogenetic tree display and annotation. Nucleic Acids Res. 49, W293–W296 (2021).

55. D. T. Jones, D. Cozzetto, DISOPRED3: precise disordered region predictions with annotated protein-binding activity. Bioinformatics 31, 857–863 (2015).

56. T. L. Bailey, J. Johnson, C. E. Grant, W. S. Noble, The MEME Suite. Nucleic Acids Res. 43, W39–49 (2015).

57. A. Drozdetskiy, C. Cole, J. Procter, G. J. Barton, JPred4: a protein secondary structure prediction server. Nucleic Acids Res. 43, W389–94 (2015).

58. A. R. Hagen, et al., In vitro assembly of diverse bacterial microcompartment shell architectures. Nano Lett. 18, 7030–7037 (2018).

59. A. Punjani, J. L. Rubinstein, D. J. Fleet, M. A. Brubaker, cryoSPARC: algorithms for rapid unsupervised cryo-EM structure determination. Nat. Methods 14, 290–296 (2017).

60. J. Zivanov, et al., New tools for automated high-resolution cryo-EM structure determination in RELION-3. eLife 7 (2018).

61. A. Rohou, N. Grigorieff, CTFFIND4: Fast and accurate defocus estimation from electron micrographs. J. Struct. Biol. 192, 216–221 (2015).

62. A. Casañal, B. Lohkamp, P. Emsley, Current developments in Coot for macromolecular model building of Electron Cryo-microscopy and Crystallographic Data. Protein Sci. 29, 1069–1078 (2020).

63. D. Liebschner, et al., Macromolecular structure determination using X-rays, neutrons and electrons: recent developments in Phenix. Acta Crystallogr. D Struct. Biol. 75, 861–877 (2019).

64. T. C. Terwilliger, S. J. Ludtke, R. J. Read, P. D. Adams, P. V. Afonine, Improvement of cryo-EM maps by density modification. Nat. Methods 17, 923–927 (2020).

65. T. C. Terwilliger, O. V. Sobolev, P. V. Afonine, P. D. Adams, Automated map sharpening by maximization of detail and connectivity. Acta Crystallogr. D Struct. Biol. 74, 545–559 (2018).

66. P. V. Afonine, et al., Real-space refinement in PHENIX for cryo-EM and crystallography. Acta Crystallogr. D Struct. Biol. 74, 531–544 (2018).

67. T. D. Goddard, et al., UCSF ChimeraX: Meeting modern challenges in visualization and analysis. Protein Sci. 27, 14–25 (2018).

68. E. Krissinel, K. Henrick, Inference of macromolecular assemblies from crystalline state. J. Mol. Biol. 372, 774–797 (2007).

